# PINPOINT: Protease INhibitor PredictiOn at the plant–pathogen INTerface using protein language models and structural modeling

**DOI:** 10.64898/2026.07.05.736646

**Authors:** Muthusaravanan Sivaramakrishnan, Balakumaran Chandrasekar

## Abstract

Cysteine and serine proteases act as an immune hub in the plant apoplast to provide robust extracellular immunity during microbial colonisation. Microbial pathogens counteract these immune proteases by inhibiting their activity using small secreted proteins (SSPs). Traditionally, SSPs with protease-inhibitory activity are predicted using sequence-dependent database searches. However, in recent years, fungal SSPs have been shown to exhibit protease-inhibitory functions despite lacking the inhibitor domain that is annotated through sequence similarity searches. Hence, a large number of these novel SSPs with putative protease inhibitor functions are missed during detection and filtered out during sequence similarity searches. This necessitates the development of newer approaches to predict SSPs lacking an annotated inhibitor domain. Machine learning approaches, such as protein language models, have emerged as powerful tools for predicting protein functions. To date, no machine learning models have been developed to predict the protease-inhibitory activities of SSPs lacking an annotated inhibitor domain. Here, we introduce a protease inhibitor prediction pipeline, PINPOINT (<u>P</u>rotease <u>IN</u>hibitor <u>P</u>redicti<u>O</u>n at plant–pathogen <u>INT</u>erface). The PINPOINT pipeline combines fine-tuned protein language model classifiers, a structure-aware autoencoder, and effector prediction into a multi-level framework for identifying SSPs with predicted protease inhibitor functions. PINPOINT predicts protease inhibitors using SSPs sequences and monomeric structures with pre-computed structures obtained from the AlphaFold Protein Structure Database or predicted using the ESMFold public API. We successfully validated the PINPOINT platform using SSPs from the plant fungal pathogen *Macrophomina phaseolina*. Notably, the PINPOINT platform robustly predicted several of these SSPs as protease inhibitors including Sequence-unrelated but structurally similar (SUSS) effectors. We further validated the inhibitory potential of these predicted *M. phaseolina* SSPs using AlphaFold Multimer (AFM) screening against candidate apoplastic soybean cysteine and serine proteases. Additionally, this platform can be used as a pre-filtering step in AFM screening approaches to reduce the number of candidates for discovering novel SSPs with protease inhibitor function for cross-kingdom plant-microbe interaction studies. The PINPOINT platform will accelerate the prediction of novel SSPs including SUSS effectors with protease inhibitor functions in proteomes of any organisms. We made the PINPOINT pipeline accessible to the research community as a web-based notebook environment for interactive computing in Google Colab, available at https://github.com/iitj-mpg-lab/PINPOINT

## Introduction

The plant apoplast is a biochemically active extracellular compartment that provides the first line of defence against invading microbial pathogens. Apoplast is enriched with a plethora of hydrolases, including proteases that contribute to immune responses (Godson and van der Hoorn, 2021). During infection, immune proteases, such as cysteine and serine proteases, accumulate in the apoplast and inhibit pathogen growth by directly targeting virulence factors or indirectly triggering immune signalling cascades (Godson & van der Hoorn, 2021; Zhang *et al*., 2025). Furthermore, these proteases act as immunity hubs by triggering a multitude of defense responses, thereby imparting resistance (Misas-Villamil *et al*., 2016; Homma *et al*., 2023a). For instance, P69B, an apoplastic tomato serine protease, acts as an immunity hub by releasing an immunogenic peptide that activates pattern-triggered immunity (Tian *et al*., 2005; Wang *et al*., 2021; Zhang *et al*., 2024) or other immune proteases such as Rcr3 (Paulus *et al*., 2020). Similarly, cysteine proteases, such as CP1 and CP2 from maize, process the phytocytokine precursor PROZIP1 in the apoplast to generate bioactive Zip1, which subsequently induces salicylic acid (SA)-dependent defense responses against biotrophic pathogens (Ziemann *et al*., 2018). For successful colonization of host plants, microbial pathogens suppress the activities of immune proteases by secreting small secretory proteins (SSPs) with >250 amino acid residues in the apoplast. SSPs belonging to diverse inhibitor families have been reported as microbial effectors that exhibit protease inhibitor functions. EPI1 and EPI10 are Kazal-like effectors secreted by *Phytophthora infestans* to inhibit the tomato P69B (Tian *et al*., 2004, 2005; Tian & Kamoun, 2005). EPIC1 and EPIC2B are cystatin-like inhibitors secreted by *Phytophthora infestans* to inhibit defence-related proteases such as C14, PIP1, and Rcr3 and to suppress the plant immune response (Jashni *et al*., 2015).

Traditionally, SSPs with protease inhibitor functions are predicted by sequence similarity searches against databases such as MEROPS or through domain searches against sequence-dependent platforms such as Interpro or Pfam. However, beyond sequence similarity, protease inhibitors possess a unique and highly conserved structural motif required for their inhibitory function (Farady and Craik, 2010; Tušar *et al*., 2021). For instance, in a recent study, AlphaFold multimer screening (AFM) of a large number of SSPs has revealed that four SSPs such as XpSSP1, CfEcp36, FoTIL, and FoSix15 from various pathogens inhibit tomato P69B (Homma *et al*., 2023b). Notably, these SSPs lacked sequence-level annotated inhibitor domains and were structurally distinct. Similarly, in our recent study, K2QS10, a SSP from *M. phaseolina* with no sequence-level similarities exhibited cysteine protease inhibitory activity (Prakash *et al*., 2025). Therefore, it is evident that a large number of these novel SSPs with putative protease inhibitor function are missed and filtered out during sequence similarity searches. AFM is emerging as an important tool for discovering protease-inhibitory functions of novel SSPs from various sources. However, these analyses are computationally intensive and require substantial Graphics Processing Units (GPU) infrastructure, limiting their scalability for large-scale screening without any pre-filtering. Therefore, a computationally efficient approach that integrates sequence and structural features for inhibitor prediction would be a valuable asset for accelerating SSP prediction across various microbial pathogens in a shorter time for cross-kingdom interaction studies. Machine learning has emerged as a powerful approach to predict protein functions in the field of plant-microbial interactions. For instance, machine learning tools such as EffectorP, Fungtion, and AMAPEC use alignment-free strategies to predict microbial effectors and antimicrobial activities of fungal effector proteins, respectively (Sperschneider & Dodds, 2022; Mesny & Thomma, 2024; Li *et al*., 2024). However, to date, no machine learning models have been developed to predict the protease-inhibitory activities of SSPs lacking the annotated inhibitor domain.

Here, we introduce a machine learning platform, PINPOINT (<u>P</u>rotease <u>IN</u>hibitor <u>P</u>redicti<u>O</u>n at plant–pathogen <u>INT</u>erface) for predicting protease inhibitor function of SSPs. In our work, we fine-tuned two state-of-the-art protein language models (PLMs), ESM-2 and ProtBERT (Elnaggar *et al*., 2021; Lin *et al*., 2023) to predict protease inhibitors. We named these fine-tuned PLMs as PIP-BERT and PIPES-M, that capture functional features beyond sequence similarity and enable the prediction of distant homologues and structurally conserved inhibitors without sequence-level motifs. Furthermore, the platform combines a structure-aware autoencoder and effector prediction into a multi-level framework for robust prediction of SSPs with protease-inhibitor functions. PINPOINT predicts protease inhibitors using SSPs sequences and monomeric structures with precomputed structures obtained from the AlphaFold Protein Structure Database or predicted using the ESMFold public API. We have validated the PINPOINT platform by using SSPs from *Macrophomina phaseolina,* a plant fungal pathogen of various legume crops. The SSPs employed for validation did not show sequence similarity to any annotated protease inhibitors. Notably, the PINPOINT platform robustly predicted several of these SSPs as protease inhibitors including Sequence-unrelated but structurally similar (SUSS) effectors. The inhibitory functions of these predicted SSPs were further confirmed using Alpha-Fold Multimer (AFM) screening against soybean cysteine and serine proteases detected in apoplast. The PINPOINT platform can also can also serve as a prefiltering step in AFM screening approaches to reduce computational costs and accelerate prediction inhibitor discovery for cross-kingdom plant-microbe interaction studies. We have made the PINPOINT pipeline accessible to the research community as a web-based notebook environment for interactive computing in Google Colab, available at https://github.com/iitj-mpg-lab/PINPOINT

## Materials and Methods

### Dataset collection and curation of the training dataset

Protease inhibitor sequences were extracted from the MEROPS database (Rawlings, 2010; Rawlings *et al*., 2016), and sequences of length between 60 and 250 amino acids were filtered. In addition, protease inhibitor sequences of similar length were extracted from the UniProt database (The UniProt Consortium, 2019) by querying with search terms “(keyword: KW-0646) AND (length: [60 TO 250])”. In total, 1,18,902 sequences were collected from both sources combined. To remove redundant and highly similar sequences, after combining sequences, they were clustered using the MMseqs2 clustering suite (--min-seq-id 0.3, -c 0.8, -s 7.5, --cov-mode 0) (Steinegger & Söding, 2017). Furthermore, sequences containing nonstandard amino acids were removed, yielding a non-redundant, homology-reduced positive dataset of 7,281 PI proteins. For negative dataset preparation, we collected 4,58,645 small secretory fungal proteins from the UniProt database by querying the search terms (taxonomy_id:4751) AND (length: [60 TO 250]) and considered only proteins with signal peptides. These sequences were clustered using the same MMseq2 threshold used for the inhibitor dataset, resulting in a 1,53,149 homology-reduced, non-redundant sequence set. The proteins lacking Pfam domain annotations were identified using Pfamscan (https://github.com/aziele/pfam_scan) with default settings and removed to ensure the dataset contains only proteins with known functions (Finn, 2005). The protease inhibitor-associated Pfam domains were identified from sequences belonging to GO terms (GO:0097179 and GO:0030414) using Pfamscan, and any sequences containing these domains were removed from the negative dataset (Binns *et al*., 2009). Additionally, the sequences showing similarity to protease inhibitors in the MEROPS and UniProt databases were identified using MMseq2 with an E-value threshold of 0.001 and removed. Finally, nonstandard amino acids in protein sequences were removed, yielding a negative dataset of 15,464 proteins unlikely to function as protease inhibitors.

### Feature descriptors extraction

The amino acid composition (AAC) and physicochemical properties between PI and non-protease inhibitor groups were calculated using the Pfeature package (Pande *et al*., 2023). The differences in AAC and physicochemical properties between protease inhibitors and non-protease inhibitors were estimated with Welch t-tests, two-sided Mann-Whitney U tests, and Benjamin-Hochberg FDR for multiple testing. The enrichment or depletion of a group in PI was assessed based on adjusted q-value (<0.05) and effect size (Cohen’s d ≥ 0.5). The protein descriptors for the protease inhibitors and non-protease inhibitors datasets were extracted using the ProtFeat package (https://github.com/gozsari/ProtFeat) (Wang *et al*., 2017; Chen *et al*., 2018). The feature descriptors were grouped into 4 hybrid descriptor sets based on their properties as follows: i) Composition-based descriptors set consists of amino acid composition, dipeptide composition, grouped amino acid composition, grouped dipeptide composition, and pseudo-amino acid composition. This feature set captures the global protein representation by encoding amino acid fractions and distributions to obtain insights into compositional differences, local residue-pair information, and sequence-order effects. ii) Autocorrelation feature descriptor set includes Moran, Geary, Normalised Moreau-Broto correlation methods that capture correlation and distribution of Physicochemical properties along the various lengths of protein reflecting functional constraints (Xi *et al*., 2010). iii) CTD feature descriptor captures global physicochemical and structural properties from protein sequences by providing a holistic representation of property composition, adjacency transitions, and positional distribution (Govindan & Nair, 2011). iv) The conjoint triad captures local interaction patterns in the sequences by considering the sequence as an overlapping triad based on physicochemical properties.

### Tree-based ensemble models

Tree-based ensemble models have demonstrated excellent predictive performance across multiple protein prediction tasks and have been shown to be superior to support vector machines, decision trees, and naive Bayes (Uddin & Lu, 2024). We used tree-based ensembles Random Forest (RF), eXtreme Gradient Boosting (XGBoost), and Light Gradient Boosting Machine (LightGBM) as classifiers implemented in scikit-learn (Pedregosa *et al*., 2011), xgboost (Chen *et al*., 2015), and lightgbm (Ke *et al*., 2017) libraries to predict protease inhibitors. The models were trained and evaluated using four descriptor sets described in the “Feature descriptors extraction” section as inputs. We used cost-sensitive learning approach to handle class imbalance (Feng *et al*., 2020). For the RF model, we used the class weight parameter class_weight = ”balanced”, and for XGBoost and LightGBM, we used the scale_pos_weight parameter to handle class imbalance. We trained the models with 80% of the training data (n=18,101) and evaluated them using a 5-fold cross-validation (CV) and an independent holdout dataset.

### Meta ensemble classifier

The Meta ensemble classifier predicts protease inhibitors by integrating gradient-boosting (XGBoost, LightGBM) and bagging (RF) models into a unified framework. The integration of RF, XGBoost, and LightGBM was implemented using a stacked generalisation ensemble approach to maximise predictive capability (Sivaramakrishnan *et al*., 2022; Wang *et al*., 2025). In this approach, RF, XGBoost, and LightGBM were used as heterogeneous base learners, trained on the input data, and their output probabilities were passed to a Logistic regression meta-learner for final predictions. This framework approximates a convex combination of non-linear decision functions, where the meta-learner learns optimal weights for probability outputs from the base models. In both base and meta learners, class imbalance was handled using cost-sensitive learning approach. We trained the models on 80% of the training data (n=18,101) and evaluated them using a 5-fold CV and an independent holdout dataset.

### Tabular foundation model

The Tabular Prior-data Fitted Network (TabPFN) is a pretrained deep learning classifier designed for tabular data types (Hollmann *et al*., 2025). TabPFN relies on prior knowledge learned from a large collection of synthetic tabular datasets and is robust to class imbalance and hyperparameter tuning (Liu & Ye, 2025; Nawaz *et al*., 2025). The tabPFN-2.5 model was implemented using scikit-learn and the tabpfn library (https://github.com/PriorLabs/TabPFN). In this work, we used the recently released TabPFN-2.5 version (Grinsztajn *et al*., 2026). We fitted the model with 80% of the training data (n=18,101), evaluated it using 5-fold CV and an independent holdout dataset.

### Large language model classifiers

The BERTforsequenceclassification and ESMforsequenceclassification architectures from Hugging Face transformers library (Wolf, Debut, Sanh, Chaumond, Delangue, Moi, Cistac, Rault, Louf, Funtowicz, *et al*., 2020) were finetuned and used as Large Language Model (LLM) classifiers for the protease inhibitors prediction task. Both models were designed for task-specific supervised finetuning of base BERT and ESM variants, and subsequent downstream classification tasks. In this study, we finetuned three base LLMs, BioBERT-v1.1, BioBERT-v1.2, and ProtBERT, from the BERT family, and three base models, esm2_t6_8M_UR50D (esm2_t6), esm2_t12_35M_UR50D (esm2_t12), and esm2_t30_150M_UR50D (esm2_t30), from the ESM2 family (Lin *et al*., 2023). The Biobert-v1.1 and Biobert-v1.2 models were BERT variants pretrained on large-scale biomedical corpora, whereas ProtBERT, esm2_t6, esm2_t12, and esm2_t30 were domain-specific models trained on protein sequence databases. We fine-tuned these models in a supervised manner by tokenising sequences with a maximum length of 250 aa, a batch size of 16, and the AdamW optimiser with a learning rate of 1e^-6^ and weight decay of 0.01. A linear learning rate scheduler without warmup was applied, and gradient norms were clipped. The class imbalance problem was addressed in the training dataset by applying class-weighted cross entropy loss with class weights determined using the inverse frequency of class labels in the training set. All model parameters were updated end-to-end during fine-tuning. The fine-tuned models were evaluated on the independent held-out set. The prediction probabilities for each class were obtained by applying the Softmax activation to the output logits. The finetuning was performed on a Linux system with an Intel Xeon Silver 4210R CPU, 64 GB RAM, and an NVIDIA RTX A5000 GPU with 24 GB vRAM. The finetuned LLM classifiers were evaluated on an independent holdout dataset. The top ranked classifiers were further evaluated on an unseen out-of-distribution (OOD) dataset to test their generalisation slightly beyond the training dataset. The positive set of the ODD dataset consists of homology-reduced protease inhibitor sequences, slightly longer than those in the training dataset curated from the Peptipedia, MEROPS, and UniProt databases. The negative set of the ODD dataset consists of proteins unlikely to have protease inhibitor activity, collected from the UniProt database, and spans three distinct protein families (kinases, DNA-binding proteins, and endopeptidases) across both viridiplantae and fungal taxa.

### Model training and evaluation

All model performance was assessed using scikit-learn.metrics module and compared in terms of metrics such as Accuracy, Precision score, recall score, F1score, Matthew’s correlation coefficient (MCC), area under the ROC curve (AUROC), and area under the precision-recall curve (AUPRC) as follows,

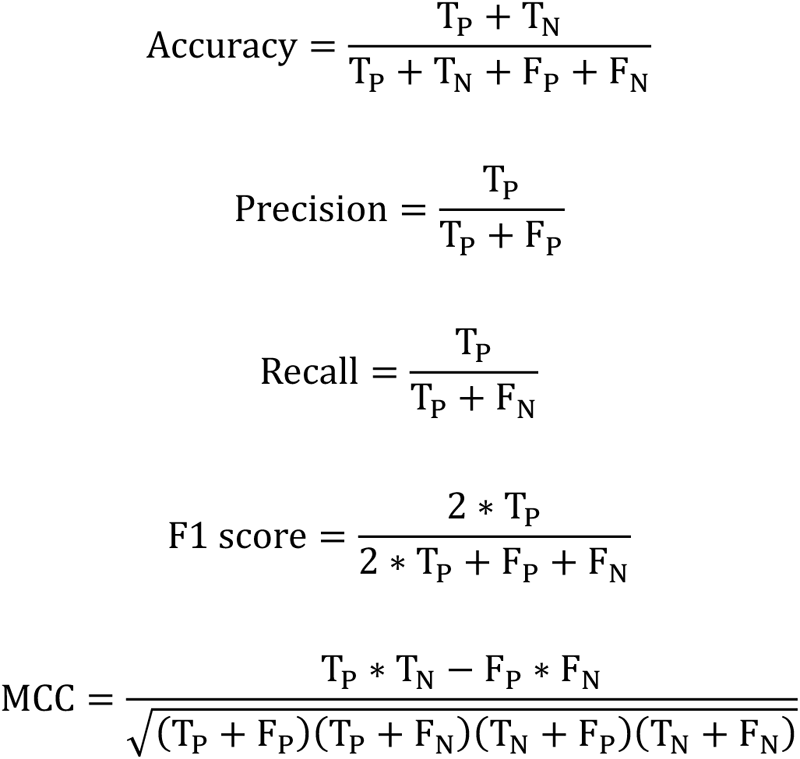

where T_P_, T_N_, F_P_, and F_N_ represent true positive, true negative, false positive, and false negative instances, respectively.

### t-distributed stochastic neighbour embedding (t-SNE) visualisation

The protein embeddings were extracted from the last layer of each pretrained and fine-tuned protein language model using attention-masked mean pooling across residues. The PCA and t-SNE analyses were performed using the Scikit-learn package. To reduce the high-dimensional noise, the embeddings were first projected into principal components using PCA (n=50). The reduced representations were then embedded into two dimensions using t-distributed stochastic neighbour embedding (t-SNE) with perplexity = 40, PCA-based initialisation, learning rate = ‘auto’, and a fixed random seed for reproducibility. To quantify cluster separability, silhouette scores were computed on the PCA-reduced embeddings, with higher values indicating improved intra-class cohesion and inter-class separation. The kernel density estimation contours were used to visualise class-specific clusters.

### Interpretability analysis

The feature attribution was calculated using the integrated gradient method implemented in the captum library (Kokhlikyan *et al*., 2020; Miglani *et al*., 2023). For each sequence, attribution scores were calculated separately for the positive (protease inhibitors) and negative (non-protease inhibitors) classes using 25 integration steps, and then normalised using a min-max scaler. Residues with attribution scores above the 75^th^ percentile were classified as high-attribution positions. A discriminative importance score was calculated by subtracting the normalised attribution score of the negative class (non-protease inhibitors) from the normalised positive class (protease inhibitors) attribution score for each residue.

### Structure-aware model

We mapped the Merops database against the UniProt database with taxonomic restriction to viridiplantae, fungi, and bacterial taxa, identified 19,973 identical hits, and downloaded their AF2-predicted structures from AFDB using the public web API (Varadi *et al*., 2024). The structural embeddings were computed for the protease inhibitor structures using the RCSB embedding model, which converts residue-level embeddings from the ESM-3 model into structure-level embeddings using a transformer-based aggregator network (Segura *et al*., 2026). Non-finite and NaN values were replaced with 0, and Z-score normalisation was applied using a standard scaler. The pre-processed embeddings were used to train a one-class unsupervised deep autoencoder implemented in the PyOD package (Chen *et al*., 2025). The Merops protease inhibitor embeddings were treated as an inlier distribution and split into training (90%) and 10 % for validation set during hyperparameter tuning. Hyperparameters of the models were tuned using Bayesian optimisation implemented in the Optuna package (Akiba *et al*., 2019). A deep autoencoder was implemented with geometric scaling in the hidden layers to ensure smooth compression. The hyperparameters were dynamically tuned for the search space bottleneck width (128–320), activation functions (ReLU, Tanh, Leaky ReLU, Sigmoid), learning rate (10−4 to 5×10−3), and dropout rates (0.1–0.4) with the objective of reducing the Mean square error (MSE) on the validation set.

### Secretome prediction

The putative secreted proteins were annotated using a predict_secretome pipeline (https://github.com/fmaguire/predict_secretome/) in permissive mode, which uses SignalP (Petersen *et al*., 2011), TMHMM (Krogh *et al*., 2001), Targetp (Almagro Armenteros *et al*., 2019), WoLF PSORT (Horton *et al*., 2007). The predicted secreted proteins with lengths < 250 aa were considered small secretory proteins (SSPs). The SSPs with known Pfam domains were identified using Pfamscan and removed. All uncharacterized SSPs were modelled using AlphaFold2 (AF2) in monomer mode with MMseqs2-generated MSA, template enabled, dropout enabled, and 20 recycles (Jumper *et al*., 2021; Mirdita *et al*., 2022). All the models were relaxed using the Amber force field.

### AlphaFold2 prediction and AFM Screening

The *M.phaseolina* SSPs were modelled with host proteases as heterodimers using AlphaFold multimer mode implemented in CUDA-supported localcolab v1.5.5 (https://github.com/YoshitakaMo/localcolabfold) (Evans *et al*., 2021; Mirdita *et al*., 2022). The localcolabfold uses Multiple sequence alignment (MSA) and PDB template obtained from the MMseqs2 search, and PDB100, respectively. The AFM screening was performed using single, with the following parameter settings: template usage enabled, dropout enabled, 20 recycles, an early recycle stop tolerance of 0.5 to control the convergence criterion, and the top tanked model was energy-minimised using the Amber force field. All runs were performed on a Linux system with an Intel Xeon Silver 4210R CPU, 64 GB RAM, and an NVIDIA RTX A5000 GPU with 24 GB vRAM. The AFM modelled SSP-protease complexes were evaluated for quality using performance metrics such as predicted template modelling (pTM) score, predicted local distance difference test (pLDDT), and interface pTM score (ipTM). The AFM confidence scores were calculated as [(0.8 × ipTM + 0.2 × pTM)] and was used as a key metric for screening protein complexes as described previously (Prakash *et al*., 2025). The spatial information about interface contacts and accuracy metrics at residue levels was evaluated using the Colabfold Batch AlphaFold-2 multimer structure analysis pipeline (Schmid, 2023). The SSP-protease complexes were visualised using Pymol (DeLano, 2002) and ChimeraX (Pettersen *et al*., 2021). The catalytic triad of the host proteases were annotated using Interpro database or MSA search. The SSPs interact within an 8Å distance of the catalytic triad were identified using NeighbourSearch algorithms of the Biopython library (Cock *et al*., 2009).

### Structural homolog identification

The *M.phaseolina* SSPs having a similar fold to known protease inhibitors were identified using foldseek structural similarity search (Van Kempen *et al*., 2024). The SSPs with TM score ≥ 0.5 were considered as structural homologs.

## Results and discussion

### Establishing a systematic machine learning (ML) framework for protease inhibitor prediction

Traditional sequence-based approaches for functional annotation often fail when a protein lacks conserved sequence motifs or sequence similarities. Nevertheless, machine learning (ML) models have been demonstrated to predict protein function using mathematical and computational representations beyond sequence alignment or homology searches. However, ML models have been reported for predicting protease inhibitors. Therefore, we aimed to develop a robust ML model for the prediction of protease inhibitors. To construct a reliable prediction model, we first curated a dataset comprising 7,281 protease inhibitor proteins and 15,464 non-protease inhibitor proteins, each 60–250 amino acid residues in length, from public sequence repositories to serve as non-redundant, homology-reduced training data for the models. The protease inhibitor set consisted of experimentally validated or expert-curated protease inhibitor sequences from the MEROPS and UniProt databases, covering all major kingdoms of life. The non-protease inhibitor set consists of carefully curated secreted proteins that are unlikely to exhibit protease-inhibitory activity. Next, we performed comparative analyses of amino acid composition (AAC) and physicochemical properties (PCP) between the curated protease inhibitor and non-protease inhibitor datasets to determine whether they exhibit distinct features. Interestingly, comparative analysis revealed statistically significant differences in AAC and PCP between protease inhibitors and non-protease inhibitor proteins (Figure S1a and Figure S1b). Amino acid residues, such as cysteine, glutamate, and lysine, were enriched, whereas alanine and leucine residues were significantly depleted in the protease inhibitor datasets. (adjusted q-value < 0.05, Cohen’s d ≥ 0.5) (Figure. S1a). The PCP features solvent accessibility (exposed), hydrophilicity, and negatively charged residues were enriched, whereas PCP features aliphaticity, hydrophobicity, neutral charge, and solvent accessibility (buried) were depleted in the protease inhibitor sequences (adjusted q-value < 0.05, Cohen’s d ≥ 0.5) (Figure S1b). Collectively, the distinct AAC and PCP profiles observed between the curated protease and non-protease inhibitor data further support the suitability of the prepared dataset as high-quality training data for establishing ML models for protease inhibitor prediction.

To develop a ML model for predicting small secreted proteins (SSPs) with protease inhibitor functions, we systematically trained and evaluated two complementary ML frameworks: a protein descriptor-based approach and a large language model (LLM)-based approach (Figure 1). In the descriptor-based approach, protein sequences are transformed into a predefined numerical representation that encodes physicochemical properties, compositional patterns, and sequence-order characteristics. Whereas, the LLM-based frameworks learn contextual sequence representations directly from large-scale text corpora using transformer architectures. For the protein descriptor-based approach, we evaluated multiple tree ensemble models such as Random Forest (RF), Extreme Gradient Boosting (XGBoost), and Light Gradient Boosting Machine (LightGBM), a stacked meta-ensemble framework, and a transformer-based tabular foundation model for predicting protease inhibitors (Figure 1a). These models were trained on four hybrid protein descriptor sets generated from the curated protease inhibitor and non-protease inhibitor datasets using the ProtFeat package. Protease inhibitor and non-protease sequences were split into training (80%) and independent holdout (20%) datasets, while maintaining class proportions. The training dataset was used to train the ML models, whereas the independent holdout dataset, which was not used during model training and reserved for robust unbiased performance evaluation to assess the generalisation ability of the trained models. Each model was systematically evaluated across descriptor types to ensure that the assessment was not biased towards any specific descriptor type or ML model using a stratified five-fold cross validation (CV) on the training dataset, followed by testing on an independent holdout dataset. For the LLM-based approach, we evaluated six fine-tuned LLMs for protease inhibitor prediction, including three ESM2 variants (esm2_t6, esm2_t12, and esm2_t30) and three BERT-family models (BioBERT-v1.1, BioBERT-v1.2, and ProtBERT). ESM2 variants and the BERT-family model, ProtBERT are domain-specific LLMs pretrained on large protein sequence databases and designed to capture the evolutionary and biochemical features encoded in protein sequences. Whereas the BERT-family models: BioBERT-v1.1 and BioBERT-v1.2 were pretrained on large biomedical text corpora, including research articles and medical literature. We evaluated these diverse sets of protein descriptor- and a large language model (LLM)-based approaches to investigate how different pretraining objectives, transformer architectures, and training data influence the performance of protease inhibitor prediction.

**Figure 1.**
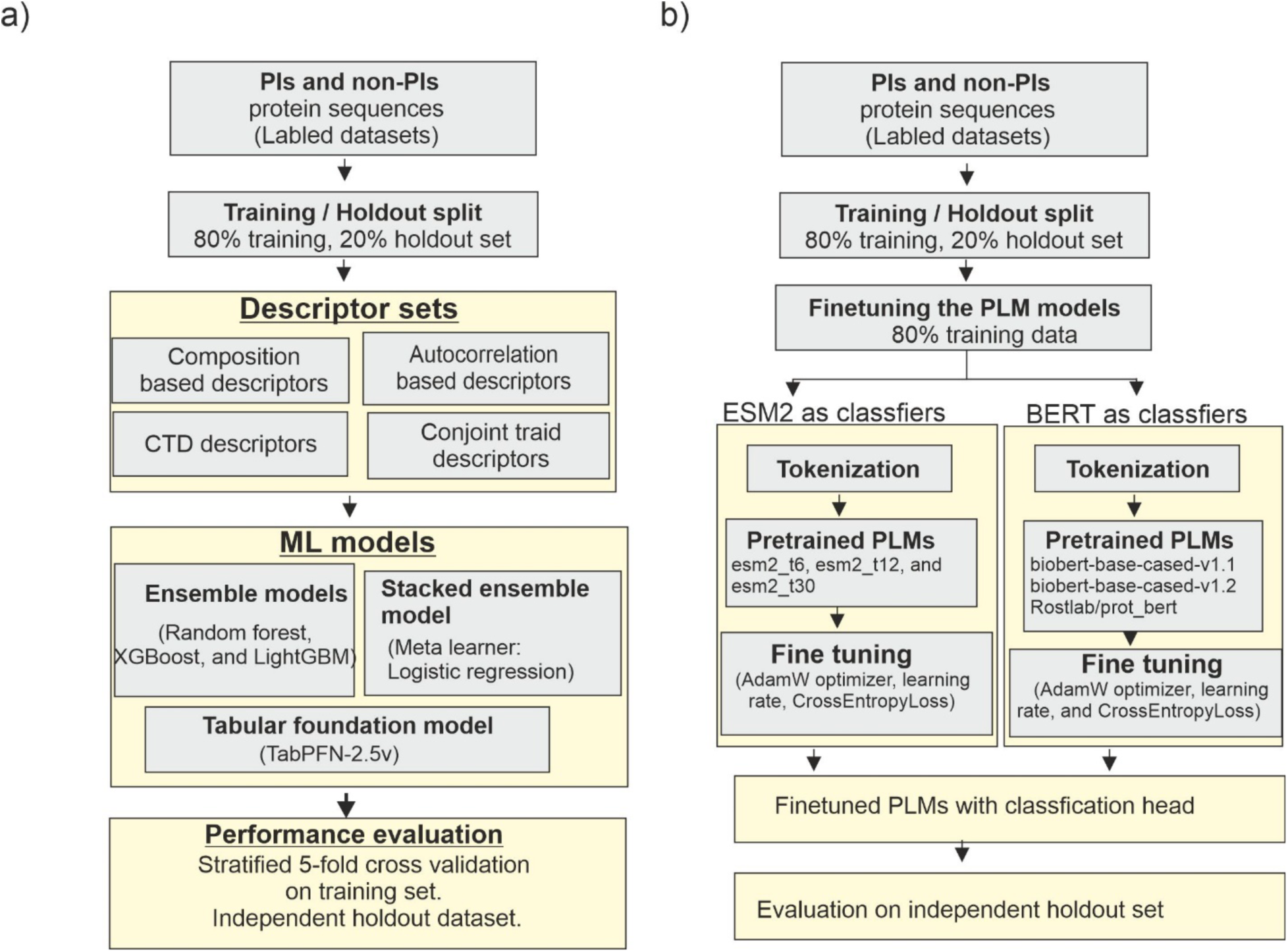
Establishing a systematic ML framework for PI prediction task. a) The workflow for systematic evaluation of a protein descriptor-based approach for the PI prediction task. PI and non-PI datasets were curated from the MEROPS and UniProt databases, clustered with the MMseq2 clustering suite to eliminate redundant sequences, and split into 80% for training and 20% for a holdout set for independent testing. Twelve protein feature descriptors were calculated for both the PI and PI datasets using the ProtFeat package. These feature descriptors were grouped into four sets for evaluation purposes as follows: (i) composition-based descriptors (amino acid composition, dipeptide composition, grouped amino acid composition, grouped dipeptide composition, and pseudo-amino acid composition), (ii) autocorrelation-based descriptors (Moran, Geary, and Normalised Moreau-Broto correlation methods), (iii) CTD descriptors (CTD Composition (CTDC), CTD Transition (CTDT), and CTD Distribution (CTDD)), and (iv) conjoint triad descriptors. These descriptor sets are used for training various ML models, including tree-based ensemble models (Random Forest, XGBoost, and LightGBM), a meta-ensemble model, and a tabular-based model (TabPFN-2.5v). The Performance of each model was assessed using a stratified five-fold CV on the training set and an independent holdout set. b) The workflow for predicting protease inhibitors using pretrained LLMs as classifiers. The LLM-based approach uses the same training and holdout sets as the protein descriptor-based approach. Both ESM2 variants (esm2_t6, esm2_t12, and esm2_t30) and three BERT-family models (BioBERT-v1.1, BioBERT-v1.2, and ProtBERT) were finetuned for the protease inhibitors prediction task with 80% training set. All the models were equipped with a linear classification head and finetuned using the AdamW optimiser and a class-weighted cross-entropy loss to address class imbalance. Finally, the predictive capabilities of these models were assessed on an independent holdout set. ML, machine learning; PI, protease inhibitor; CTD, Composition, Transition, and Distribution; CV, cross-validation; ESM2, Evolutionary Scale Modelling 2; LightGBM, Light Gradient Boosting Machine; XGBoost, Extreme Gradient Boosting. TabPFN, Tabular Prior-Data Fitted Network; LLMs, large language models; BERT, Bidirectional Encoder Representations from Transformers; BioBERT, Biomedical Bidirectional Encoder Representations from Transformers; PLM, pretrained language model; ProtBERT, Protein Bidirectional Encoder Representations from Transformers.

### The tabular foundation model outperforms ensemble and metaensemble models on the protease inhibitor prediction

To evaluate the predictive performance of the descriptor-based approach, we compared the ensemble, metaensemble, and TabPFN-2.5v models across four descriptor sets using an independent holdout dataset (Figure 2). The ensemble models combine predictions from multiple base learners to improve accuracy and robustness by reducing prediction errors (Jain *et al*., 2026). Metaensemble models further integrate multiple model outputs through a meta-learner to enhance predictive performance (Harun-Or-Roshid & Kurata, 2025). TabPFN-2.5 is a transformer-based foundation model for tabular data that performs in-context learning, enabling accurate predictions (Liu & Ye, 2025; Grinsztajn *et al*., 2026). F1-score was used as the primary ranking metric in the analysis, as it provides a balanced measure of precision and recall, thereby reflects the overall performance of the classifiers (Christen *et al*., 2024). MCC and ROC-AUC were used as complementary metrics of imbalance-aware performance and overall class discrimination, respectively. For ensemble and meta-ensemble models, we observed comparable performance between the CV and independent holdout datasets across all tested descriptor sets, indicating strong generalisation (Table S1 and Table S2). The composition-based descriptors demonstrated moderate predictive performance across all classifiers, whereas the tree-based ensemble models performed consistently well (Figure 2a). The XGBoost delivered the best results using the composition-based descriptor set, achieving the highest test accuracy (0.8959), F1-score (0.8349), ROC-AUC (0.9535), and MCC (0.7589). For all these models evaluated, autocorrelation and conjoint triad features obtained lower AUROC scores, suggesting that these descriptors have weaker discriminative power for classifying protease inhibitors from non-protease inhibitors (Figure 2b and Figure 2c). Interestingly, the Composition, Transition, and Distribution (CTD) feature set consistently yields the best results across all evaluated classifiers. By integrating information on amino acid composition, sequence order, and residue distribution, the CTD feature set provides a robust representation of protein sequences, which likely contributes to its consistent superior performance. Among the ensemble models assessed, XGBoost demonstrated the strongest overall performance with CTD descriptors, achieving an F1-score of 0.8676, MCC of 0.808, and a ROC-AUC of 0.9677 on the independent holdout dataset. (Figure 2d). Using CTD descriptors, the metaensemble outperformed XGBoost with an F1-score of 0.8709, MCC of 0.8134, and ROC-AUC of 0.9693 on the independent dataset (Figure 2e). Although the improvement was modest, the consistent increase in MCC, together with the maintained balance between precision and recall, suggests that the meta-ensemble offers a more robust and reliable prediction than the ensemble models. Notably, the TabPFN-2.5v consistently outperformed both ensemble and meta-ensemble models across all evaluated descriptor sets (Table S3). Among all model descriptor sets tested, TabPFN-2.5v with CTD descriptors achieved the highest overall predictive performance, with an accuracy of 0.9381, precision of 0.9248, recall of 0.8758, F1-score of 0.8996, and log-loss of 0.1598 on the independent dataset (Figure 2f). Furthermore, despite the strong predictive performance of descriptor-based models, none of the tested models exceeded an F1 score of 0.90 (Figure 2g). Achieving an F1 score of more than 0.90 would provide a more reliable platform for the protease inhibitor prediction with fewer errors. Collectively, these analyses suggest that protein descriptor-based ML approaches may not adequately capture the sequence characteristics associated with protease inhibitors, prompting us to evaluate LLM-based approaches.

**Figure 2.**
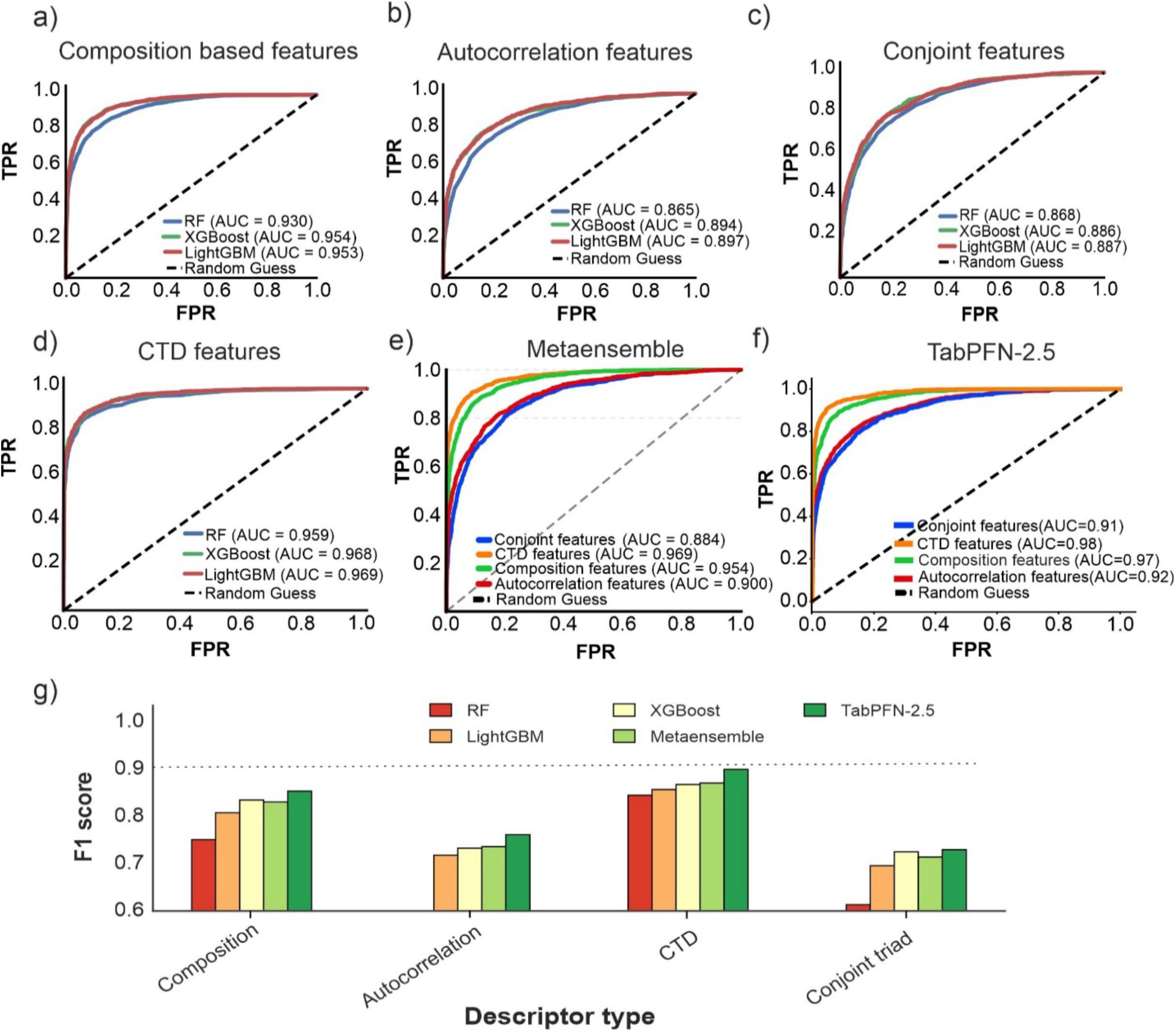
The tabular foundation model outperforms ensemble and metaensemble models on the PI prediction task. The ROC curves of tree-based ensemble models across four distinct feature descriptor sets: (a) composition-based descriptors, (b) autocorrelation-based descriptors, (c) conjoint triad descriptors, and (d) CTD descriptors. (e) ROC curves demonstrating the predictive performance of the meta-ensemble model on four feature descriptors evaluated. (f) ROC curves of the tabular foundation model (TabPFN-2.5v) on four feature descriptors evaluated. All models were assessed using both a five-fold stratified CV and an independent test set. (g) The classification performance of tree-based ensembles, the meta-ensemble, and the tabular foundation model (TabPFN-2.5v) on the independent holdout set was ranked by F1 score. The dotted horizontal line indicates the threshold value of F1 score greater than 0.90, which is the target value needed for an ML model to minimise false positive predictions. PI, protease inhibitor; ROC, receiver operating characteristic; CTD, Composition, Transition, and Distribution; TabPFN, Tabular Prior-Data Fitted Network; CV, cross-validation; ML, machine learning.

### Finetuned protein language models achieve state-of-the-art classification performance on the protease inhibitor prediction

To overcome the limitations of the protein descriptor-based approach, we next investigated whether task-specific fine-tuned LLMs could improve protease inhibitor prediction. Task-specific finetuned LLMs and their embeddings have recently been shown to perform well on protein function prediction and classification (Schmirler *et al*., 2024; Yuan *et al*., 2025). We fine-tuned six LLMs with training data and evaluated them on an independent test set using the same metrics as the descriptor-based approach. For model training, we used protease inhibitor and non-protease inhibitor sequences of lengths ranging from 60 to 250 amino acid residues. Therefore, during inference on longer input queries (> 250 aa), the first 250 amino acid residues from the N-terminus were extracted and used for analysis. The ProtBERT and ESM2 models pretrained on protein sequence databases consistently outperformed general biomedical language models (BioBERT-v1.1 and BioBERT-v1.2) on the protease inhibitor prediction (Table S4). The finetuned ProtBERT (30 layers, 420M parameters) model achieved the best overall performance, with an F1-score of 0.9847, MCC of 0.9776, and ROC-AUC of 0.999 (Figure 3a). Interestingly, ESM2 variants (esm2_t6 and esm2_t12) also demonstrated reasonable predictive capability, but their performance is clearly lower than that of the esm2_t30 model (30 layers, 150M parameters), indicating that predictive capability increases with model size (Table S4). Among the tested ESM2 models, the fine-tuned esm2_t30 model achieved the highest performance, with an F1-score of 0.9666, MCC of 0.9514, and ROC-AUC of 0.9931 (Figure 3b). The best performing fine-tuned PLMs, ProtBERT and esm2_t30, we hereafter referred as PIP-BERT and PIPES-M, respectively.

**Figure 3.**
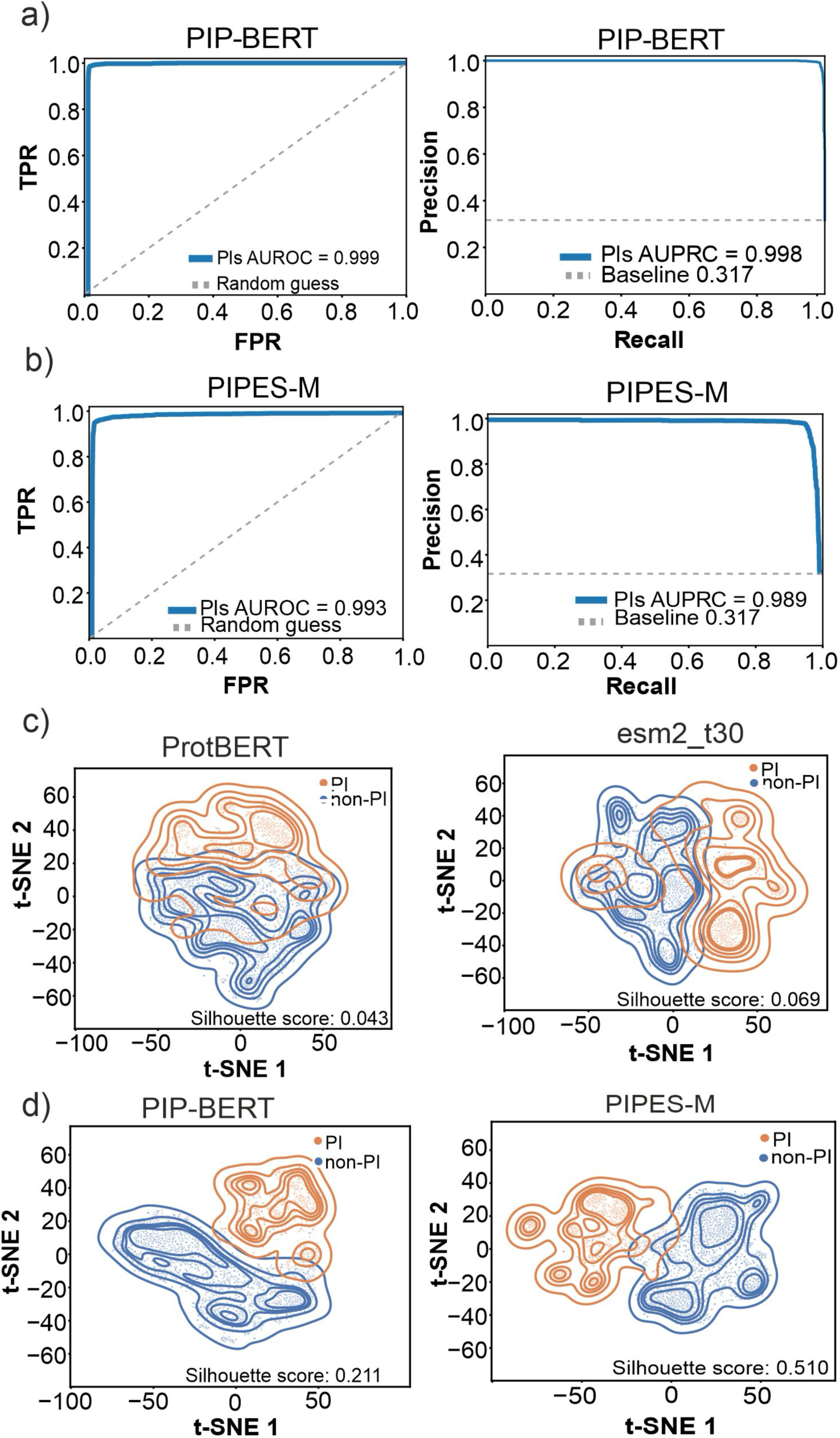
Fine-tuned PLMs achieve state-of-the-art classification performance on the protease inhibitor (PI) prediction task. **(a,b)** ROC and PR curves for the best performing fine-tuned protein language models, PIP-BERT (a) and PIPES-M (b), evaluated on the independent test set. High AUROC and AUPRC scores for the finetuned PLMs indicate state-of-the-art classification performance on the PI prediction task. In the ROC plots, the dashed diagonal line denotes a random guess (AUROC = 0.5). The ROC curves for both models remain substantially above this reference line, indicating excellent discrimination between PI and non-protease-inhibitor (non-PI) sequences. In the PR plots, the dashed horizontal line represents the baseline prevalence of the positive class. The PR curves for both models remain well above this baseline, demonstrating high precision across a broad range of recall values and highlighting their ability to accurately identify PI sequences while minimising false positives. (c) The t-SNE visualisation of embeddings extracted using pretrained ProtBERT and esm2_t30 models before finetuning, showing substantial overlap between PI and non-PI sequences and low silhouette scores. (d) t-SNE visualisation of embeddings extracted using the corresponding fine-tuned models, PIP-BERT and PIPES-M, showing compact and distinctly separated PI and non-PI clusters with increased silhouette scores. The protein embedding space is represented as a density contour and coloured by class label. PLMs, protein language models; PI, protease inhibitor; non-PI, non-protease inhibitor; ROC, receiver operating characteristic; PR, precision–recall; AUROC, Area Under the Receiver Operating Characteristic curve; AUPRC, Area Under the Precision–Recall curve; t-SNE, t-Distributed Stochastic Neighbour Embedding; ProtBERT, Protein Bidirectional Encoder Representations from Transformers; esm2_t30, Evolutionary Scale Modelling 2 model with 30 transformer layers.

To complement the metrics-based assessment of the finetuned PLMs, we visualised protein embedding space using t-Distributed Stochastic Neighbour Embedding (t-SNE). t-SNE is a technique for visualising high-dimensional feature spaces in 2D, whereas the silhouette score quantifies the degree of cluster separation. Herein, t-SNE analysis was used to investigate how protease inhibitor and non-protease inhibitor classes are organised in the latent space before and after fine-tuning. This analysis provides visual evidence that fine-tuning reshapes the latent embedding space by incorporating task-specific supervisory signals to make protease inhibitor and non-protease inhibitor classes linearly separable. Before fine-tuning, pretrained PLMs form heavily overlapping clusters between protease inhibitor and non-protease inhibitor classes, as evidenced by low silhouette scores (Figure 3c). This highlights the limited capacity of PLMs to adequately represent features relevant to specialised functional tasks without supervised finetuning. Meanwhile, both fine-tuned models, PIPES-M and PIP-PERT, form compact, distinctly separated clusters for protease inhibitor and non-protease inhibitor classes, as further evidenced by a substantial increase in silhouette scores (Figure 3d). This further demonstrates that task-specific fine-tuning restructures the embedding space to represent features related to protease inhibitors, thereby improving the ability to identify and distinguish them from non-protease inhibitors. In addition, to further assess the added value of end-to-end fine-tuning over pretrained PLMs for protease inhibitor prediction, we performed an ablation analysis. In this analysis, for each LLM, the pretrained backbone was frozen, and only the classification head was trained using the same input dataset and parameters used for end-to-end fine-tuning. Notably, all ablation models demonstrated a noticeable decrease in predictive performance compared to their fine-tuned versions (Table S5). The noticeable difference in predictive capability between fine-tuned and ablation models shows that the robust performance of PIPES-M and PIP-BERT results from end-to-end fine-tuning rather than from inherent pretraining knowledge. Taken together, these results indicate that model size and domain-specific pretraining are critical factors in determining the predictive capability of LLM-based protease inhibitor predictions. Further, fine-tuned PLMs PIP-BERT and PIPES-M outperforms the descriptor-based model for protease inhibitor predictions.

### PIP-BERT and PIPES-M models show robust prediction on sequences of different lengths and low sequence identity regimes

Although PIP-BERT and PIPES-M models achieved high predictive performance on the independent test set, it is crucial to verify whether they generalise well across distant homologs and varying sequence length distributions. We performed additional robustness analyses on an independent holdout dataset stratified by sequence length (60–250 amino acid residues) and pairwise sequence identity (30–80%). Interestingly, both PIPES-M and PIP-PERT models maintained high performance even at the low sequence identity bins (30–40% and 40-50%), indicating strong generalisation to remote homologs (Figure 4a and Table S6). The stable performance of the fine-tuned PLM models across low sequence identity bins suggests effective learning of functional signals beyond sequence similarity. Similarly, both PLMs demonstrated consistently high predictive performance across all sequence length bins tested (F1 > 0.94; ROC-AUC > 0.98) (Figure 4b). The PIP-BERT exhibited length-independent generalisation with minimal performance variation (F1 > 0.97; ROC-AUC 0.99) across all ranges (Figure 4b and Table S7). The PIPES-M showed robust performance (F1: 0.943-0.975), with a slight reduction in predictive capability for the length range 141-220 amino acid residues (Table S7 and Figure 4b). Collectively, these results indicate that the fine-tuned PIP-BERT and PIPES-M models generalise effectively across both sequence length and low sequence identity regimes, making them suitable for predicting novel protease inhibitors from protein sequences lacking conserved domains or detectable homologs.

**Figure 4.**
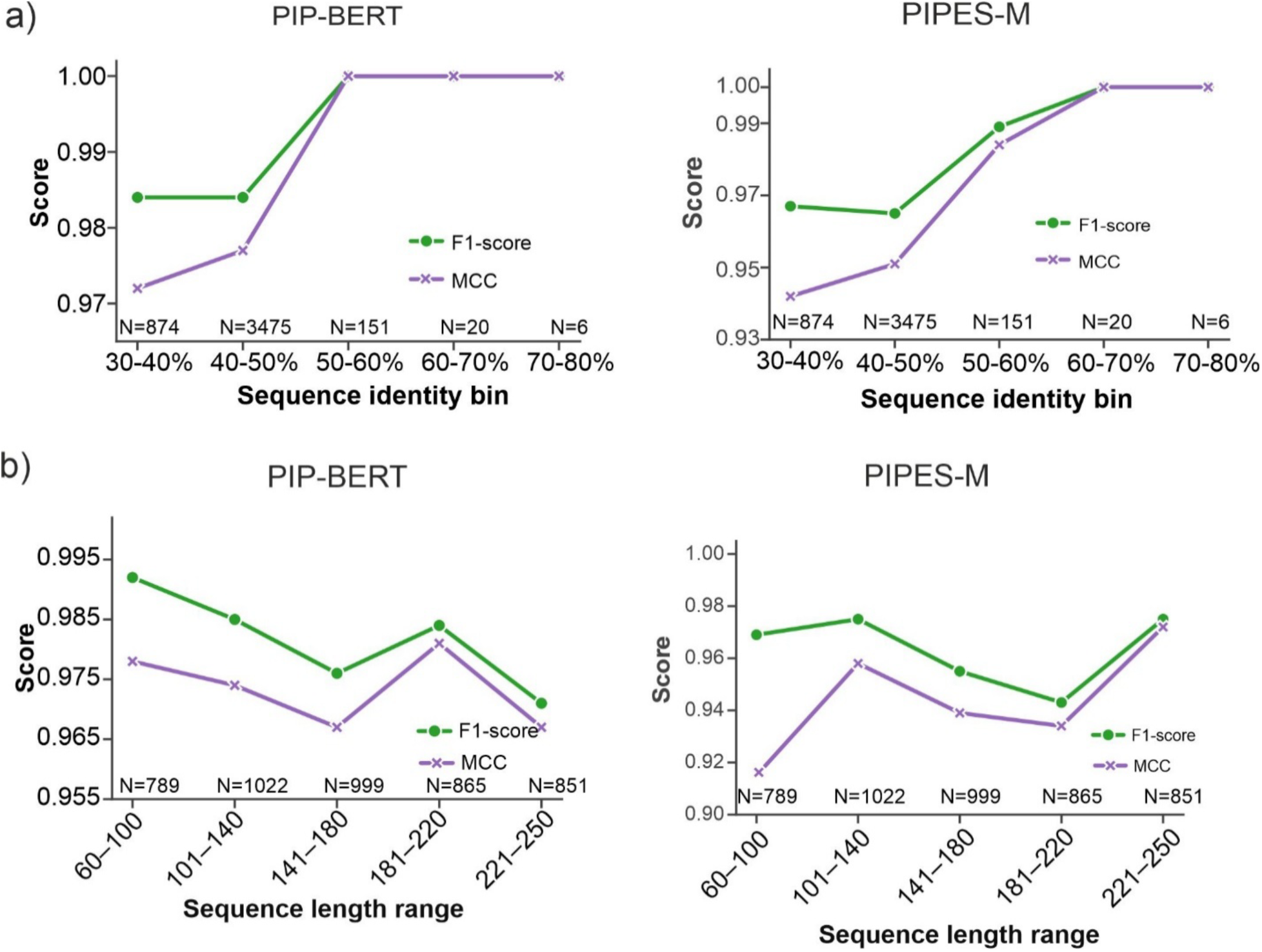
PIP-BERT and PIPES-M models demonstrate robust performance across varying sequence identity and length regimes. **(a)** The performance of the PIP-BERT and PIPES-M models was evaluated on an independent test set partitioned into pairwise sequence identity bins ranging from 30% to 80%. F1-score (green circles) and MCC score (purple crosses) remained consistently high across all identity ranges, including low sequence identity bins. The number of samples (N) within each sequence identity bin is indicated above the x-axis. (b) Performance of PIP-BERT and PIPES-M across different protein sequence length ranges. F1-score (green circles) and MCC score (purple crosses) remained consistently high across all sequence length bins, indicating that model performance is independent of protein length. MCC, Matthews correlation coefficient. N, Number of proteins in each bin.

When evaluating model performance on data drawn from the same distribution as the training set, there is a possibility of a bias toward the training data distribution and results in overestimation of models predive capability. The training dataset used for model development comprised predominantly of short protease inhibitor sequences representing all major kingdoms of life, while non-protease inhibitor sequences were restricted to fungal taxa. To investigate the extent to which the models generalise beyond on data shifts from the training distribution, we assessed PIPES-M and PIP-BERT on a non-biased out-of-distribution (OOD) dataset. In this OOD dataset, the positive set consists of protease inhibitor sequences that are slightly longer than the training data (>250 amino acid residues), and the negative set consists of proteins annotated as non-protease inhibitor from a wide-range of functional classes such as kinases, DNA-binding proteins, and endopeptidases covering viridiplantae and fungi taxa, ensuring a challenging distribution shift for stringent assessment of the models on protease inhibitory prediction task. Interestingly, both PIP-BERT and PIPES-M models displayed a high AUROC (>0.95) in the OOD dataset, highlighting their capacity to capture generalised sequence–function relationships (Figure 5a and Figure 5b). Both models demonstrated strong predictive performance for protease inhibitor versus non-protease inhibitor on the OOD dataset, indicating robust generalisation across varying lengths of protease inhibitor sequences (Figure 5c and Table S8). Furthermore, the robust classification of kinases, DNA-binding proteins, and endopeptidases from the Viridiplantae and Fungi taxa as non-protease inhibitors demonstrates strong cross-taxonomic generalisation with reduced lineage-specific bias (Figure 5d). Collectively, these findings demonstrate that both PIP-BERT and PIPES-M exhibit strong and consistent generalisation under challenging distribution shifts, including remote homology, variable sequence lengths, and cross-taxonomic out-of-distribution settings, underscoring their robustness and reliability. Taken together, these results indicate that PIP-BERT and PIPES-M exhibit robust and consistent generalisation across remote homology, variable sequence lengths, and cross-taxonomic OOD datasets, which underline their robustness and reliability.

**Figure 5.**
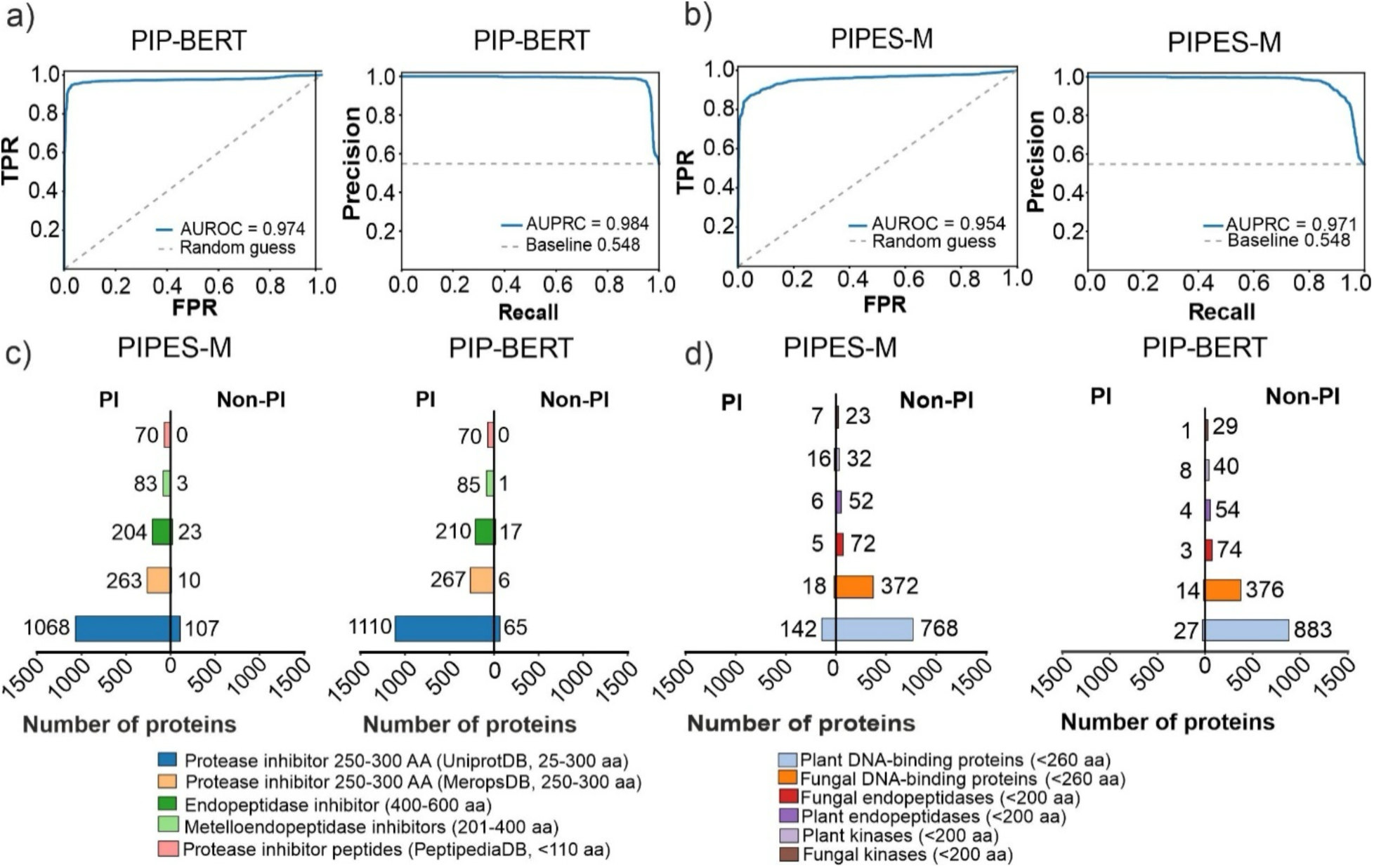
PIPES-M and PIP-BERT models show robust performance on out-of-distribution (OOD) datasets. ROC and precision–recall PR curves for PIP-BERT (a) and PIPES-M (b) models evaluated on the OOD dataset. The dashed diagonal line in the ROC plots represents a random guess (AUROC = 0.5), whereas the dashed horizontal line in the PR plots represents the baseline positive class prevalence. Both models maintain strong classification performance on the ODD dataset, with ROC and PR curves remaining substantially above their respective reference lines. (c) Performance of PIP-BERT and PIPES-M models on PI sequences slightly longer than the training dataset distribution. Most PI sequences were correctly classified across the length, demonstrating robust length extrapolation and generalisation. (d) Performance of PIP-BERT and PIPES-M on a taxonomically diverse dataset consists of non-PI proteins from fungal and viridiplantae taxa. Only a small number of non-PI sequences were misclassified as PI, indicating strong cross-taxonomic generalisation and robustness to sequence diversity. PI, protease inhibitor; non-PI, non-protease inhibitor; ROC, receiver operating characteristic.

### Interpretability analysis on experimentally verified protease inhibitors reveals attribution patterns of PIP-BERT and PIPES-M models

To investigate if the fine-tuned PIP-BERT and PIPES-M models can be employed to predict protease inhibitor function of microbial small secreted proteins (SSPs), we analysed six SSPs from various fungal pathogens, such as XpSsp1 (*Xanthomonas perforans*), CfEcp36 (*Fulvia fulva*), SDE1 (*Candidatus Liberibacter asiaticus*), FoTIL (*Fusarium oxysporum*), FoSix15 (*Fusarium oxysporum*), and K2QS10 (*Macrophomina phaseolina*). These SSPs have a sequence length of ≤250 amino acid residues and have been experimentally validated for their protease-inhibitor function, despite lacking the annotated protease-inhibitor domain (Shabab *et al*., 2008; Homma *et al*., 2023a; Prakash *et al*., 2025). Remarkably, the PIPES-M model classified all six SSPs as protease inhibitors, whereas the PIP-BERT model classified five of the six SSPs as protease inhibitors, except the K2QS10 protein (Table S9). This analysis suggests that these models differ in their attribution patterns when predicting protease inhibitor functions.

To investigate the attribution patterns of PIP-BERT and PIPES-M models, we performed an interpretability analysis using EPIC2B and K2QS10 using the integrated gradient method. The integrated gradient method assigns an importance score to each residue in a protein sequence to quantify its contribution to the classification output of the models (Sundararajan *et al*., 2017). EPIC2B is a characterised cysteine protease inhibitor with well-annotated cystatin-like domain, whereas K2QS10 is a characterised cysteine protease inhibitor which lacks the annotated domain. Residue-level raw attribution profiles revealed significant differences in the patterns for both EPIC2B and K2QS10 (Figure S2). Next, we calculated the attribution profile difference by subtracting positive (protease inhibitor) and negative (non-protease inhibitor) class attribution scores to identify the key residues contributing to protease inhibitor prediction. The positive attribute profile difference score indicates that residues preferentially support the protease inhibitor class, whereas negative scores indicate that residues support the non-protease inhibitor class. The attribution difference profiles for EPIC2B indicated a strong positive contribution from N-terminal residues to protease inhibitory activity in both the PIP-BERT and PIPES-M models (Figure 6a). This aligns with the contribution of the N-terminal tripartite wedge in EPIC2B for protease inhibitor functions (Kaschani & Van Der Hoorn, 2011). This suggests that both PIP-BERT and PIPES-M models capture conserved, functionally relevant motif-driven features in EPIC2B. Interestingly, K2QS10 exhibited a scattered attribution pattern with the PIPES-M model covering multiple secondary structure elements across the entire sequence, whereas the PIP-BERT model exhibited strong signals with negative attribution difference scores in support of the non-protease inhibitor class (Figure 6b). These contrasting attribution profiles in the case of K2QS10 suggest that PIP-BERT and PIPES-M rely on different predictive features for functional inference. To further investigate the prediction modes of the PIP-BERT and PIPES-M models, we performed self-attention map analysis using EPIC2B and K2QS10. The self-attention map analysis reveals how the model progressively transforms sequence information from local biochemical features into global structural and functional representations (Vig *et al*., 2021; Chandra *et al*., 2023). The self-attention maps were extracted from various transformer layers of the PIP-BERT and PIPES-M models and compared with the corresponding AF2 structure-derived residue-residue distance matrices for both EPIC2B and K2QS10. For EPIC2B, both the PIPES-M and PIP-BERT models capture long-range residue interaction patterns, as evidenced by the off-diagonal signals in the self-attention maps (Figure 6c). However, PIPES-M showed stronger and more pronounced off-diagonal signals across successive layers, suggesting a more robust representation of long-range residue dependencies (Figure 6c). In the case of K2QS10, PIPES-M identified long-range residue interaction patterns associated with the predicted secondary structural fold relevant to protease inhibitory function, whereas PIP-BERT failed to detect these structural features (Figure 6d). Taken together, the attribution and self-attention map analyses indicate that while both models can capture sequence-derived features for protease inhibition, PIPES-M additionally leverages long-range structural relationships that are critical for recognizing non-canonical protease inhibitors. This indicates that PIP-BERT is primarily constrained to the local sequence context, whereas PIPES-M captures both local interactions and higher-order structural topology signals.

**Figure 6.**
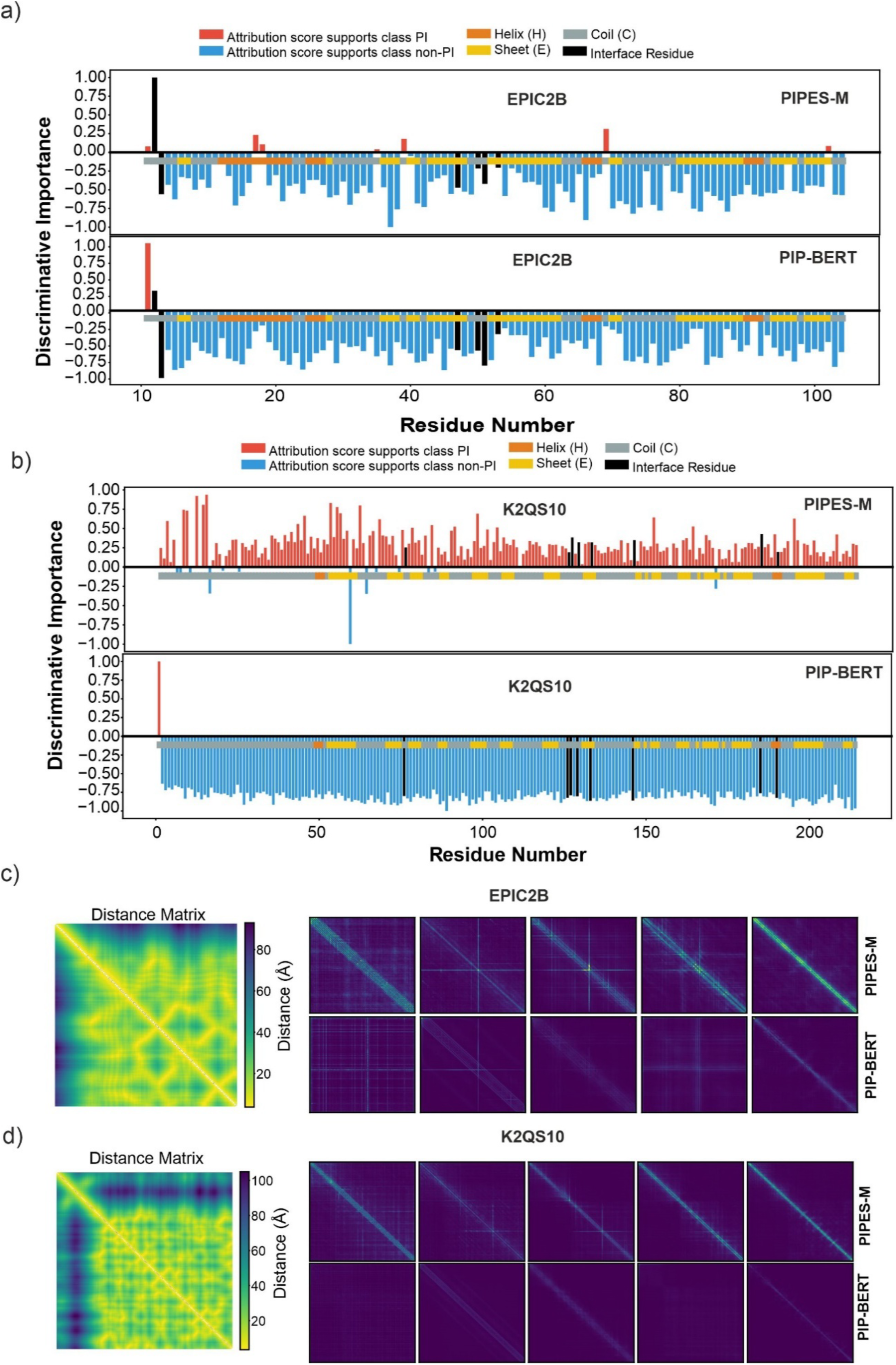
An interpretability analysis of experimentally verified Inhibitors EPIC2B and K2QS10 reveals model-specific attribution patterns. The residue-level interpretability was performed using the integrated gradients algorithm implemented in the captum library for experimentally verified inhibitors (a) EPIC2B from *P. infestans* and (b) K2QS10 from *M. phaseolina*. The discriminative importance score was calculated as the difference between positive (PI) and negative (PI) class attribution scores. The bar heights indicate the relative importance of each residue. The red bars representing attribution score support PI, and the blue bars indicate attribution score support non-PI. The black bar denotes AFM-predicted interface residues. The secondary structure elements (SSE) were calculated for both inhibitors using their PDB structures downloaded from AFDB and displayed below the attribution profiles (orange: α-helix; yellow: β-sheet; grey: random coil). The comparison of multi-head attention maps and residue-residue distance matrix for EPIC2B (c) and K2QS10 (d). The leftmost panels display the ground-truth residue-residue distance matrices (Cα–Cα distances) calculated from AF2 high-confidence modelled structures. The adjacent grids present the multi-head attention maps for PIPES-M (top row) and PIP-BERT (bottom row) across representative transformer layers (Layers: 6, 12, 18, 24, and 30). PI, protease inhibitor; non-PI, non-protease inhibitor; IG, Integrated Gradients; AFM, AlphaFold-Multimer; PLCP, papain-like cysteine protease; AFDB, AlphaFold Protein Structure Database; PDB, Protein Data Bank; SSE, secondary structure element; Cα, alpha carbon.

### PINPOINT pipeline predicts protease inhibitor function of *M. phaseolina* SSPs that lacks annotated inhibitor domain including SUSS effectors

We implemented the fine-tuned PIP-BERT and PIPES-M models to establish a platform we named PINPOINT (Protease INhibitor PredictiOn at plant–pathogen INTerface). PINPOINT is an integrated multi-level screening pipeline to systematically predict protease inhibitory function of SSPs that lacks the annotated inhibitor domain. The established PINPOINT pipeline consists of three sequential levels (Figure 7a). At level 1, we implemented the fine-tuned PIP-BERT and PIPES-M models to predict protease inhibitors from sequence queries (Figure 7a). In order to further enhance the protease inhibitor prediction, at level 2, we employed a structure-aware one-class deep autoencoder (StructAE) model to filter out candidates that lacks structural features for protease inhibitor function (Figure 7a). The StructAE model was trained on RCSB structural embeddings of protease inhibitor structures curated from AFDB (Segura *et al*., 2026). The StructAE model was further hyperparameter-tuned using a dedicated validation dataset comprising known protease inhibitors, and the optimal model was identified using Bayesian optimisation. The tuned StructAE model was subsequently evaluated using an unbiased, independent test set comprising protease inhibitor peptides from PeptipediaDB, as well as non-protease inhibitor protein classes, including endopeptidases (Plant and Fungi), kinases (plant and fungi), and chaperones (covering all taxa). The monomeric AF2 predicted protein structures of these datasets were queries to the tuned StructAE model and the reconstruction error was estimated. Interestingly, the tuned StructAE model demonstrated a clear separation between protease inhibitors and non-protease inhibitor protein classes (Figure S3). Protease inhibitor peptides from PeptipediaDB exhibited significantly lower reconstruction mean squared error (MSE) values than plant and fungal endopeptidases, kinases, and chaperones (Figure S3). This result demonstrate that StructAE effectively captures structural characteristics of protease inhibitors and can serve as a structure-guided prefiltering step for the reliable, high-throughput identification of SSPs with protease inhibitor-like structural features. At level 3, EffectorP, a machine learning tool, has been integrated into the pipeline to predict effector activities of proteins (Figure 7a).

**Figure 7.**
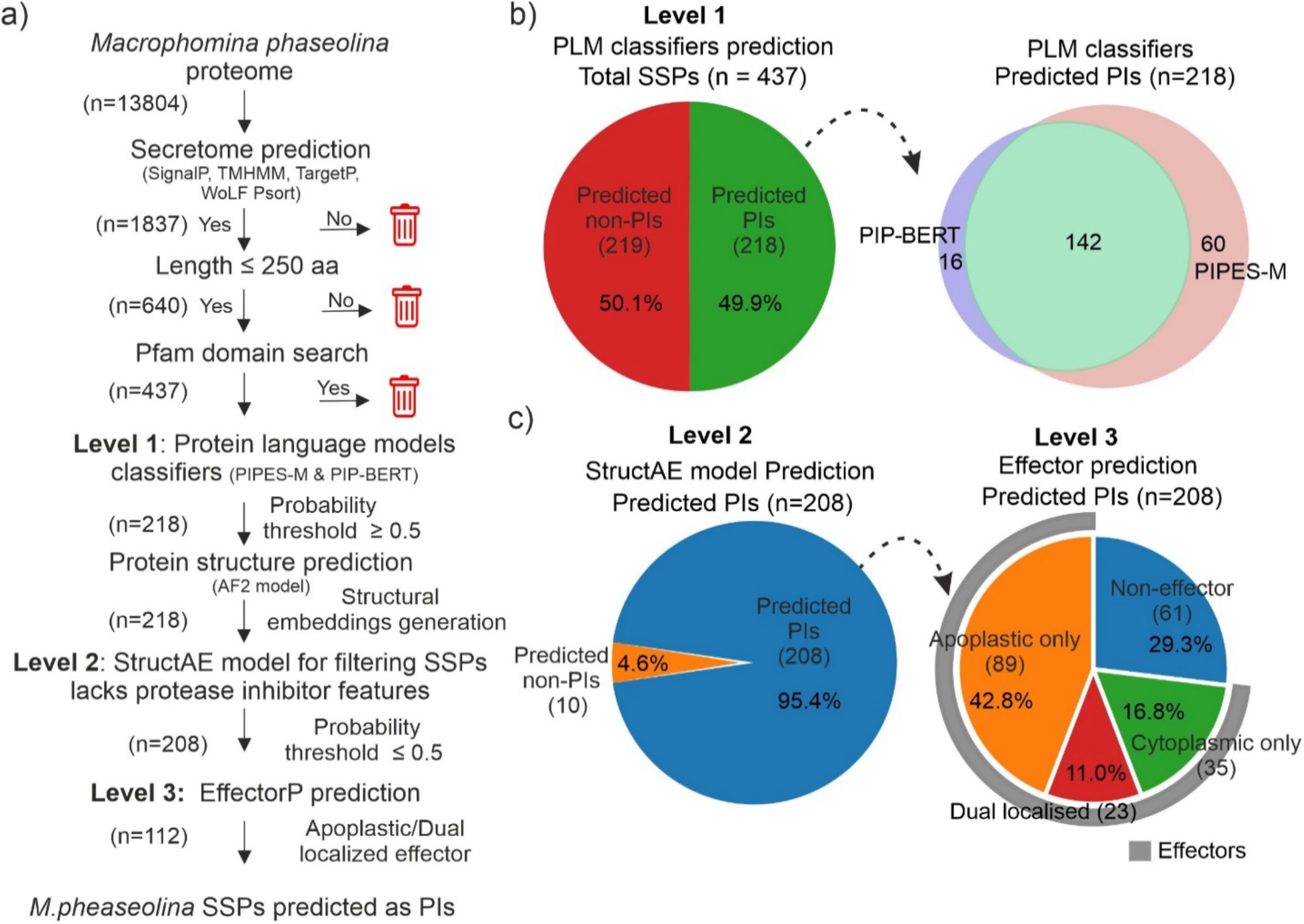
PINPOINT pipeline predicts *M. phaseolina* SSPs lacking known functional domains as potential protease inhibitors. (a) The Multilevel workflow of the PINPOINT pipeline. The secreted proteins from the *M. phaseolina* proteome were predicted using SignalP, TMHMM, TargetP, and WoLF PSORT. All proteins containing known Pfam domains or exceeding 250 amino acids in length were removed. Level 1 employs finetuned PLM classifiers (PIPES-M and PIP-BERT) to predict. Level 2 uses the StructAE model and structural embedding to filter candidates lacking PI-like features. Level 3 uses EffectorP prediction to select apoplastic/dual localised effectors among predicted protease inhibitors. (b) The *M. phaseolina* SSPs predicted as protease inhibitors by the finetuned PLM classifiers PIPES-M and PIP-BERT at Level 1. Breakdown showing the number of *M. phaseolina* SSPs predicted as protease inhibitors by PIP-BERT and PIPES-M, highlighting both unique and overlapping predictions. c) The *M.phaseolina* SSPs predicted as protease inhibitors by the StructAE model at Level 2. The breakdown of effectors secreted by *M. phaseolina* was predicted using EffectorP 3.0. Pfam, Protein families; SSPs, Small secreted proteins; PLM, Protein language models; AF2, AlphaFold2; RCSB, Research Collaboratory for Structural Bioinformatics; PDB, Protein Data Bank; TMHMM, transmembrane hidden Markov model; StructAE, Structure ware autoencoder. PI, protease inhibitor; non-PI, non-protease inhibitor;

Having established the PINPOINT pipeline, we applied the platform to predict the protease-inhibitory function of SSPs lacking the annotated inhibitor domain in *M. phaseolina*, a soil-borne plant fungal pathogen. The *M. phaseolina* genome is predicted to encode 13,804 proteins, of which 1,837 proteins were identified as secretory proteins based on signal peptides and other secretion-related features. Of these 1837 proteins, 640 proteins qualified to be annotated as SSPs based on their sequence length ≤ 250 amino acid residues. Pfam analysis of these 640 proteins has revealed that 437 SSPs did not have any known functional domain annotation (Figure 7a). The mature sequences of 437 SSPs were subjected to the established PINPOINT pipeline for predicting protease inhibitor function. At level 1, both models overlappingly predicted 142 SSPs (32.5%) as protease inhibitors, while 60 SSPs (13.7%) and 16 SSPs (3.7%) were uniquely predicted by PIPES-M and PIP-BERT, respectively (Figure 7b). Therefore, at level 1, 218 SSPs that were classified as protease inhibitors by at least one of the two fine-tuned PLM models. These candidates were subject to level 2 in the PINPOINT pipeline (Figure 7). At level 2, the 218 SSPs were modelled using the AlphaFold2 (AF2) platform and screened with the tuned StructAE model. Of the 218 candidate SSPs, 208 SSPs were robustly predicted as protease inhibitors based on structural embeddings, whereas 10 SSPs were classified as non-protease inhibitors and removed from the pipeline (Figure 7c). At Level 3, of these predicted 208 SSPs, 152 were predicted as effectors and 66 SSPs were classified as non-effectors (Figure 7c). Of these 152 predicted effectors, 89 SSPs were classified as apoplastic effectors, 35 SSPs as cytoplasmic effectors and 23 SSPs as dual-localised effectors (Figure 7c). Therefore, 112 SSPs were predicted for effector activities in the apoplast. Taken together, the established PINPOINT pipeline follows a systematic procedure to predict protease inhibitors by integrating sequence, structural, and effector predictions to systematically identify protease inhibitor–like SSPs at the proteome scale.

To validate our PINPOINT-predicted *M. phaseolina* SSPs, we employed AlphaFold Multimer (AFM) screening approach. AFM screening is an effective platform for validating the protease inhibitor-protease complexes with high confidence based and ipTM and AFM confidence scores (Bao *et al*., 2026). For the AFM screening, we employed the 112 *M. phaseolina* SSPs predicted at level 3 against the cysteine and serine proteases detected in soybean root apoplast during *M. phaseolina* infections (Prakash *et al*., 2025). We used five soybean cysteine proteases (UniProt IDs: I1M2Y6, I1JTM0, I1MER7, I1MHG0, and I1MXL3) and six soybean serine proteases (UniProt IDs: A0A0R0FVK4, I1MNL7, A0A0R0FXY0, I1MN82, I1KHC9, and I1LAF3) as baits and SSPs from *M. phaseolina* as ligands in the AFM screening. We employed our previously established AFM workflow and parameters for screening highly confident protease inhibitor-protease complexes. This workflow involved modelling soybean mature proteases and *M. phaseolina* SSPs as heterodimeric complexes using localcolabfold using MMSeqs2 search and PDB100 template followed by amber relaxation (Prakash *et al*., 2025). In parallel, we also performed the AFM screening with the *M. phaseolina* SSPs that are predicted as non-protease inhibitors by the PINPOINT pipeline. In total, 219 SSPs were predicted as non-protease inhibitors of which 43 SSPs were predicted as effectors secreted into apoplast and these SSPs were employed as a control in this analysis (Figure S4). Interestingly, of the 112 *M. phaseolina* SSPs, 20 SSPs formed a high-confidence complexes with soybean cysteine proteases and 31 SSPs formed a high-confidence complexes with soybean serine proteases with AFM confidence score ≥0.70 (Figure 8a, Figure 8b and Table S11, Table S13). In this analysis, SSPs were observed to exhibit either targeted interactions with one of the chosen soybean proteases or multiple interactions with the chosen soybean proteases (Table S10 and Table S12). For instance, five SSPs (UniProt IDs: K2S7D9, K2SEI6, K2R2M7, K2R751, and K2RSF4) were predicted to interact with more than one soybean cysteine protease and seven SSPs (UniProt IDs: K2RFU0, K2RGG5, K2RSF4, K2RYN9, K2S4X8, K2SAJ0, and K2SJ30) were predicted to target more than one soybean serine protease (Table S10 and Table S12). Furthermore, eight SSPs (K2RFU0, K2R751, K2RUD2, K2RDK5, K2RKZ2, K2RSF4, K2S4X8, and K2SYW1) displayed interactions with both soybean cysteine and serine proteases (Table S10 and Table S12). On the other hand, AFM screening of 43 SSPs predicted as non-protease inhibitors against the chosen cysteine and serine proteases revealed only three high-confidence complexes for cysteine proteases and six high-confidence complexes for serine proteases (Figure 8a and Table S11, Figure 8b and Table S13). Furthermore, we evaluated the distributions of AFM confidence scores and predicted TM (pTM) values across all modelled protease–SSP complexes. For both the cysteine and serine protease screens, SSPs predicted as protease inhibitors by the PINPOINT pipeline demonstrated significantly higher AFM confidence and pTM scores than the non-protease inhibitors (Figure S5a and Figure S5b). Interestingly, majority of the SSPs that displayed high confidence interactions with their respective proteases displayed a strong interfacial contact near the catalytic triad with the distance of around 8 Å (Table S10 and Table S12). Furthermore, the structural analysis of these complexes revealed a characteristic loop region of SSPs interacting with the catalytic site of the proteases (Figure 8c and Figure 8d). Collectively, of the 112 PINPOINT predicted *M. phaseolina* SSPs, 43 SSPs displayed strong interactions with the chosen proteases indicating their potential role as protease inhibitors. The other 69 PINPOINT predicted *M. phaseolina* SSPs that did not display interactions with our analysis might be due to the fact that these SSPs might target other families of cysteine or serine proteases or might target other classes of proteases, such as aspartate or metalloproteases, that were not used in the analysis, which requires further validation. Notably, K2QS10, a *M. phaseolina* SSP that we characterised previously as a SUSS effector, has been predicted as a protease inhibitor in the PINPOINT-pipeline and also displayed interactions with soybean cysteine protease in our AFM analysis (Table S10). This suggests that the PINPOINT-pipeline can be used to identify SUSS effectors with protease inhibitor functions. To verify this, we structurally analysed K2RFU0, a SSP, which is predicted to be a protease inhibitor and interacted with both soybean serine and cysteine proteases in our AFM screening (Figure S6a and Table S10, Table S12). Foldseek analysis against AlphaFoldDB revealed that K2RFU0 shares structural similarity with a cystatin domain-containing protease inhibitor from *Pyrus ussuriensis × Pyrus communis* (A0A5N5GTM8), with a significant TM-score of 0.603 and an RMSD of 2.43 Å (Figure S6b). Interestingly, multiple sequence alignment of these proteins shows only 18% sequence identity between them (Figure S6c). Therefore, the established PINPOINT pipeline also predicts SUSS effectors that have putative protease-inhibitor functions. Taken together, these results demonstrate that the established PINPOINT platform predicts protease inhibitor functions in SSPs lacking the annotated inhibitor domain including the SUSS effectors. Furthermore, the PINPOINT platform can also be integrated as a pre-filtering step to enrich candidates in AFM screening of protease-inhibitor complexes. This could facilitate the discovery of novel SSPs in an accelerated manner with limited computational infrastructures.

**Figure 8.**
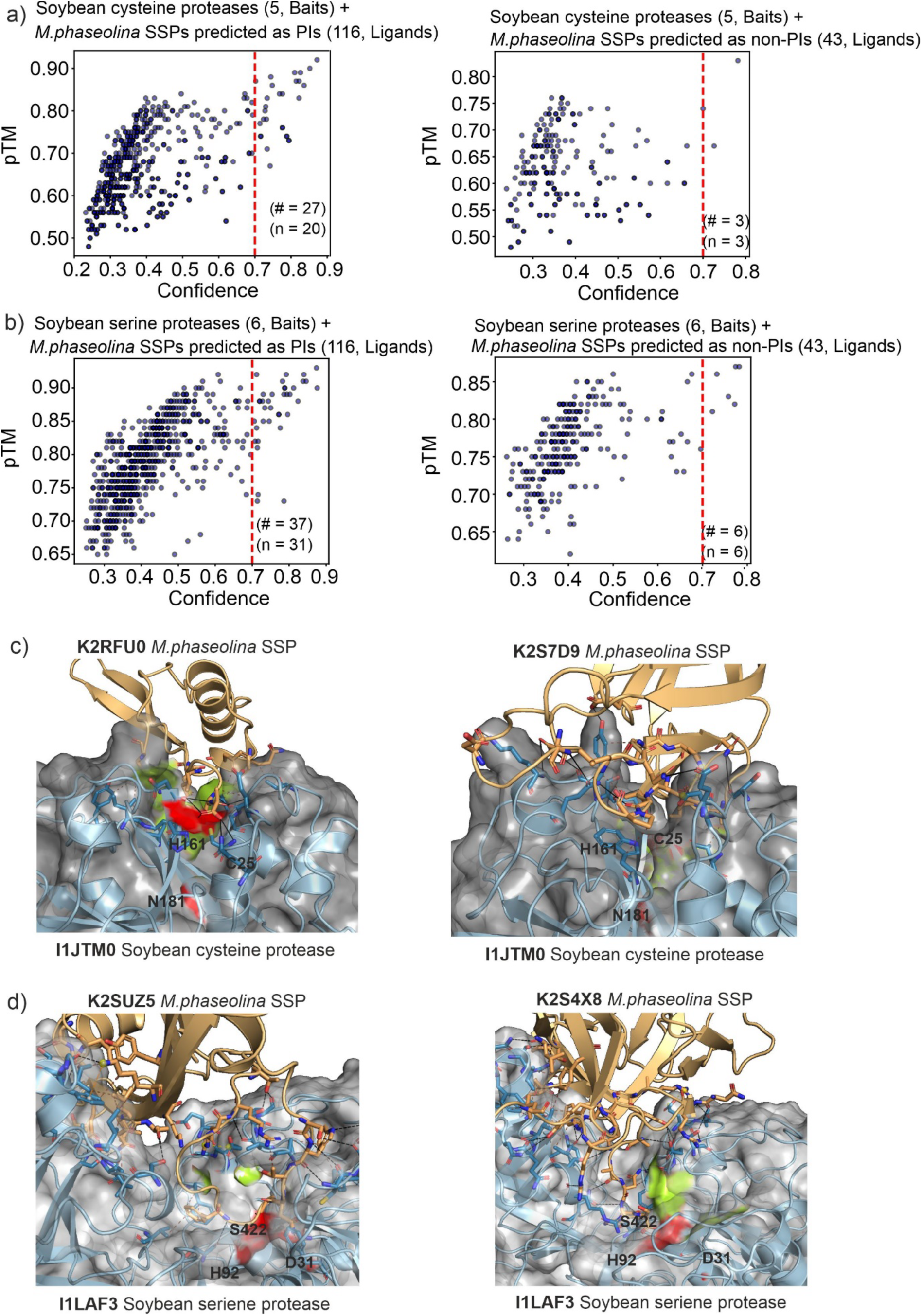
AFM screening reveals *M.phaseolina* SSPs predicted as PI by the PINPOINT pipeline form a higher frequency of high confidence complexes with soybean proteases than SSPs predicted as non-protease inhibitors. (a) The *M. phaseolina SSPs* predicted as protease inhibitors and non-protease inhibitors were screened against five soybean cysteine proteases and six serine proteases detected in the root apoplast. This analysis was implemented in the local ColabFold pipeline using the AFM confidence score as a decision metric. AFM models with a confidence score above 0.7 are considered high-confidence complexes. *M. phaseolina* SSPs predicted as PI by the PINPOINT pipeline form larger subpopulation-high-confidence complexes with AFM confidence scores > 0.7 (AF confidence scores = 0.2 * pTM + 0.8 * ipTM). (c) *M. phaseolina* SSPs (Uniprot IDs: K2RFU0 and K2S7D9) predicted as protease inhibitors by the PINPOINT pipeline interact with the active site of the soybean cysteine protease (Uniprot ID: 1JTM0). The AFM-predicted interactions were visualised using the open-source software PyMOL. The zoomed-in view of the heterodimeric interface between the *M. phaseolina* SSPs and the soybean cysteine proteases is presented. The soybean cysteine protease is displayed as a light-blue cartoon with a grey surface representation, while the *M. phaseolina* SSPs are shown as wheat-coloured cartoons. The active site is predicted using InterProScan and highlighted in green, while the catalytic triad residues are shown as red sticks and labelled with single-letter amino acid codes. (d) AFM predicted structural models of the candidate *M. phaseolina* SSPs (Uniprot IDs: K2SUZ5 and K2S4X8) interacting with soybean serine protease (Uniprot ID: I1LAF3). The soybean serine protease is represented as a light-blue cartoon with a grey surface, and SSPs are shown in wheat-colored cartoon representation. InterProScan predicted active site residues are highlighted in green, while catalytic triad residues are shown as red sticks. # = no. of confident complexes and n = no. of SSPs forming the confident complexes.

## An online prediction platform for protease inhibitors using PINPOINT

To streamline the PINPOINT pipeline and improve its accessibility to the scientific community, we have developed an online platform to access it. We implemented an easy-to-use user interface for model inference on the Google Colab platform, providing a readily accessible environment for users to predict protease inhibitors without dedicated local GPU infrastructure or complex software installation. The pipeline implemented in Google Colab consists of multiple sequential notebooks to progressively filter potential protease inhibitor candidates, as detailed in the GitHub repository, and is openly available at https://github.com/iitj-mpg-lab/PINPOINT. The PINPOINT pipeline can be executed stepwise using Google Colab notebooks as follows: First, run the “Sequence Module” Colab notebook to process protein sequences using the fine-tuned PIPES-M and PIP-BERT models. Next, for candidates predicted as protease inhibitors by the sequence module, run the “AlphaFold Fetch” notebook to retrieve AF2-predicted protein structures from AFDB using UniProt IDs as input. For novel or newly added sequences not available in AFDB, users can use the “ESMFold API” notebook, which uses the official ESMFold web API to predict 3D structures. Subsequently, use the “Structural Module” notebook to filter candidates using the StructAE model and retain proteins with PI-like structural features. This is followed by the “EffectorP Module” notebook, which uses the EffectorP tool to predict effector proteins. Finally, SSP predicted as effectors can be modelled as heterodimeric complexes with target proteases using the AFM structure prediction platform (https://github.com/google-deepmind/alphafold). All steps except AFM modelling can be performed directly on the Google Colab free-tier runtime.

## Conclusion

In this study, we systematically evaluated conventional descriptor-based machine learning approaches and fine-tuned PLMs for protease inhibitor prediction. Through rigorous evaluation on the independent test set, we demonstrated that finetuned PLMs PIP-BERT and PIPES-M achieved state-of-the-art performance and maintained strong predictive capability across low-sequence-identity and variable-length. The fine-tuned PLMs can robustly classify protease inhibitors from non-inhibitor protein classes curated from plants and fungi taxa. The fine-tuned PLMs were integrated with a structure-aware autoencoder trained on protein structural embeddings and an effector prediction module to implement the multi-level PINPOINT pipeline. The PIPOINT pipeline predicted several *M. phaseolina* SSPs that lacked the annotated protease inhibitor-associated domain as protease inhibitors including SUSS effectors. We validated the PINPOINT pipeline using AFM, and the AFM screening demonstrated that *M. phaseolina* SSPs predicted as protease inhibitors by the PINPOINT pipeline exhibit a significantly higher propensity to form high-confidence heterodimer complexes with soybean cysteine and serine proteases. These findings demonstrate that the PINPOINT can identify novel SSPs with protease inhibitor functions despite lacking the sequence homology. AFM is a crucial tool for predicting protease inhibitor interactions at the plant–pathogen interface, but it is computationally expensive, requires dedicated storage and GPU infrastructure for large-scale screening. Therefore, a pre-filtering step is necessary to reduce the candidate search space by prioritising proteins with relevant functional annotations, enabling efficient large-scale screening (Xia *et al*., 2023). Moreover, recently, the scale of human interactome prediction using AFM has been considerably reduced with the KIRC (Knowledge Integrated Rapid Classifier) model (Schmid *et al*., 2025). Similarly, the PINPOINT pipeline can be used as an effective pre-filter for AFM screening to identify novel protease inhibitors and prioritise them for AFM, thereby substantially reducing computational time, GPU resource requirements, and data storage demands. A key limitation of the PINPOINT pipeline is that it is designed for SSPs with a maximum sequence length of 250 amino acid residues. The current pipeline does not supports the prediction of proteins that are larger than 250 amino acid residues. Further versions with datasets of protein sequences with larger amino acid sequences can be trained using the LLMs employed in the PINPOINT pipeline to facilitate this prediction. To facilitate PINPOINT use by the research community, we have made it publicly available via Google Colab notebooks, providing an accessible platform for protease inhibitor prediction without dedicated computational infrastructure or complex software installation.

## Author contributions

MS: conceptualization and design, performed the bioinformatics analyses, generated figures and visualizations, and GitHub repository, writing – original draft, review and editing. BC: conceptualization and design, funding acquisition, project administration, resources, supervision, writing – original draft, review and editing.

## Supporting information

All supplemental tables

## Acknowledgements

This study work was supported by SRG grant (SRG/2022/000528) and ARG grant (ANRF/ARG/2025/003442/LS) provided by Anusandhan National Research Foundation (ANRF) (formerly Science and Engineering Research Board (SERB) India). We thank BITS Pilani, Pilani Campus, for providing the Institute Fellowship to Muthusaravanan Sivaramakrishnan. We thank Prof. Kumaravel Kandaswamy, Centre for Excellence in Microscopy, Department of Biotechnology, Kumaraguru College of Technology, for kindly providing GPU access to carry out this work.

## Data Availability Statement

The PINPOINT pipeline is openly available on GitHub repository (https://github.com/iitj-mpg-lab/PINPOINT), and the data generated during the analysis is available in the Zenodo repository (https://zenodo.org/records/20552909). The fine-tuned model weights and architecture configuration generated during this study are openly available in the Hugging Face repository at (https://huggingface.co/MuthuS97) under the repository ID (MuthuS97/PIPES-M,MuthuS97/PIP-BERT, and MuthuS97/StructAE)

## Supplementary figures

**Figure S1.**
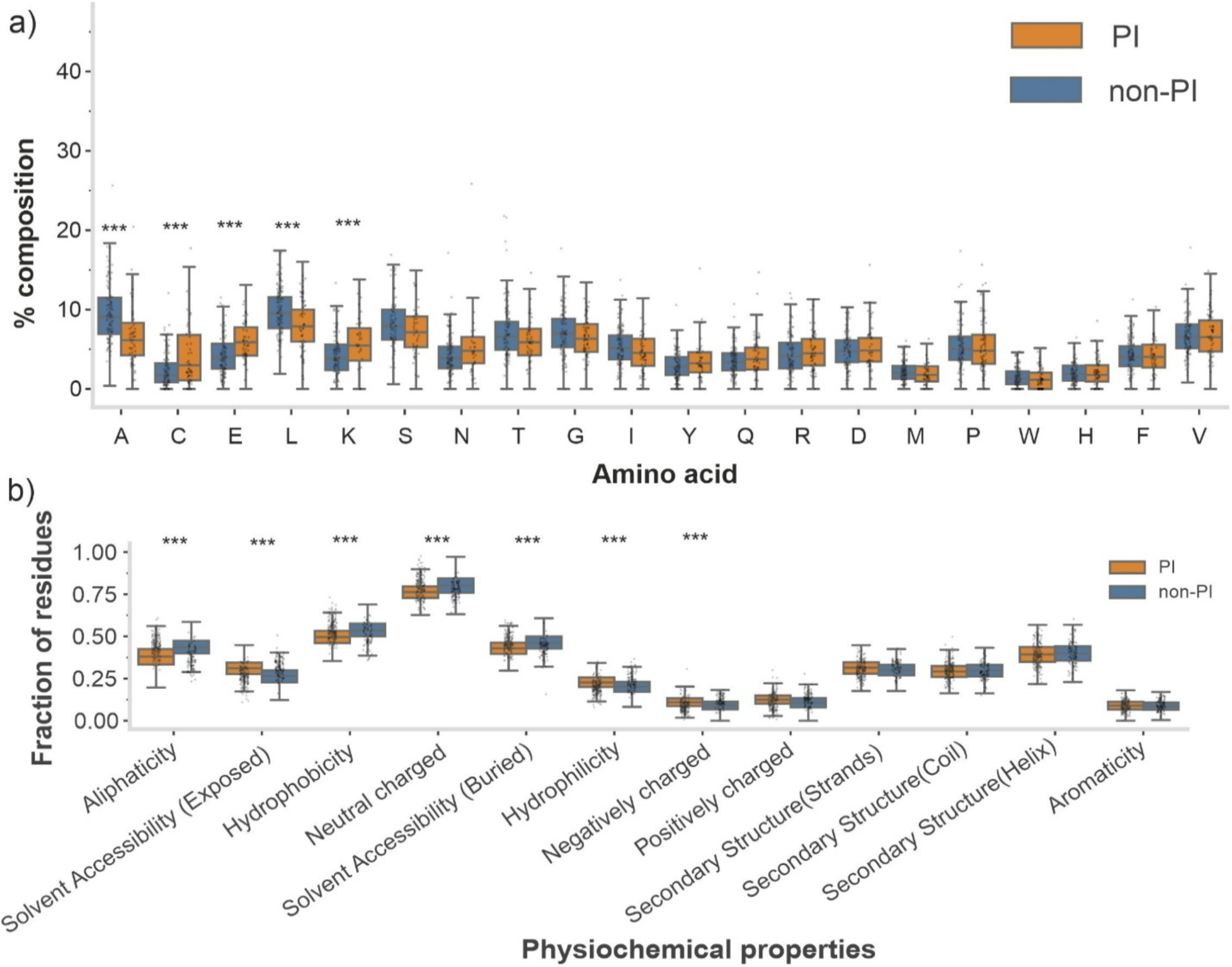
Protease inhibitors exhibit distinct sequence and physicochemical signatures. **(a)** The comparison of the amino acid composition (AAC) between PI and non-PI sequences. Box plots representing the percentage composition of 20 standard amino acids. **(b)** Comparison of the physicochemical properties (PCPs) between protease inhibitor and non-protease inhibitor sequences. Each amino acid and PCP property was compared using a two-sided Welch’s t-test, with multiple-testing correction applied using the false discovery rate (FDR) method to obtain q-values. Statistical significance was further evaluated using a two-sided Mann–Whitney U test, and effect sizes were quantified using Cohen’s *d.* The AAC and PCPs were considered significantly enriched or depleted when q < 0.05 and |Cohen’s d| > 0.5. Statistical significance is denoted as * (q < 0.05), ** (q < 0.01), and *** (q < 0.001). PI, protease inhibitor; non-PI, non-protease inhibitor;

**Figure S2.**
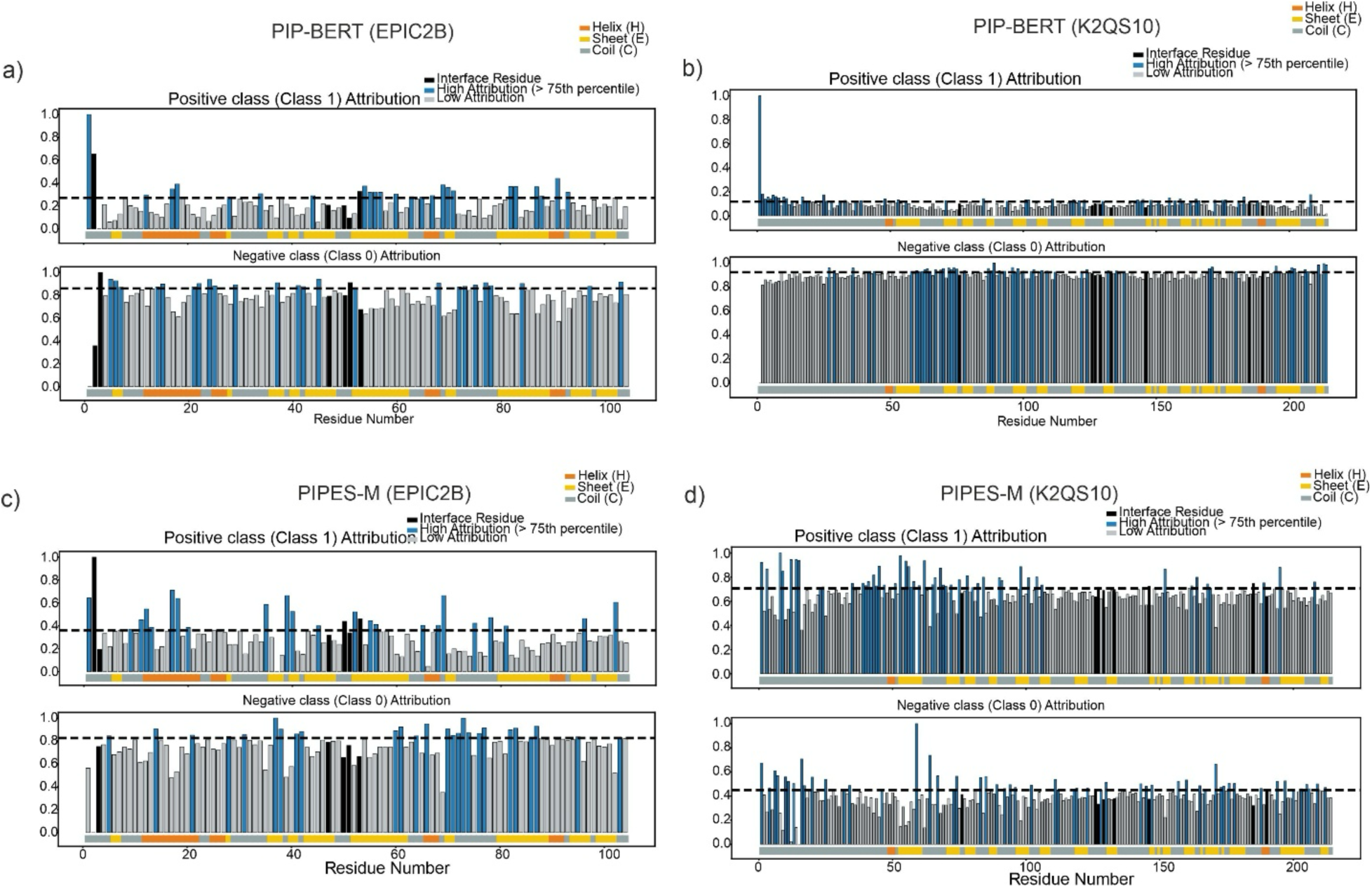
Residue-level raw attribution profiles revealed notable differences in the patterns for both EPIC2B and K2QS10. Raw Integrated Gradients (IG) attribution for EPIC2B (a, c) and K2QS10 (b, d). The dashed horizontal line represents the percentile-based threshold (75th percentile) used to identify high-attribution residues. IG, Integrated Gradients; PI, Protease Inhibitor; PLM, Protein Language Model; SSE, Secondary Structure Elements. The black bar denotes AFM-predicted interface residues. The secondary structure elements (SSE) derived from PDB structures are mapped below the profiles (orange: α-helix; yellow: β-sheet; grey: random coil).

**Figure S3.**
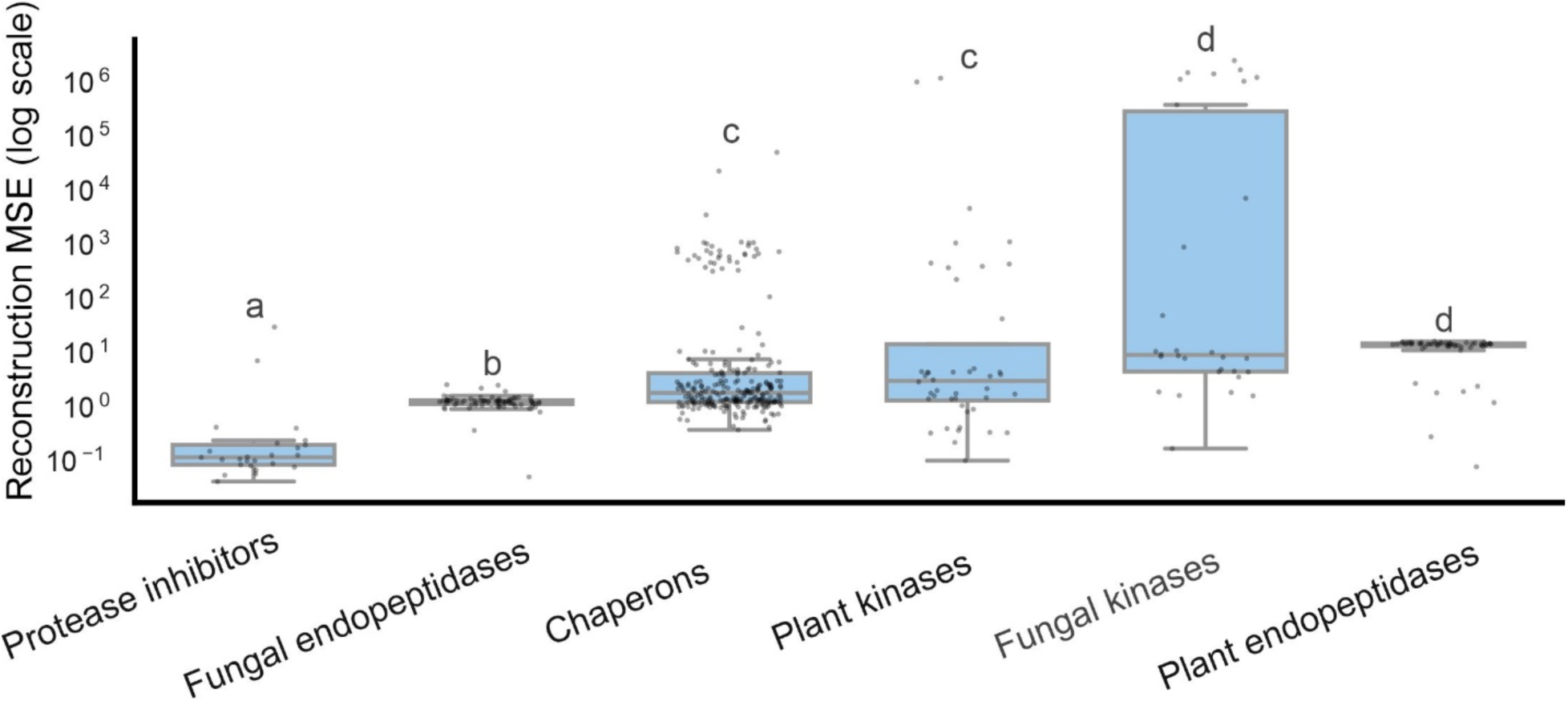
The structure-aware autoencoder (StructAE) model distinguishes protease inhibitors from non-protease inhibitor protein classes. Reconstruction mean squared error (MSE) values for protease inhibitors were compared with those of five non-protease inhibitor protein families to evaluate the ability of the StructAE model to discriminate between protease inhibitors and non-protease inhibitor proteins. Boxplots show the median (centre line) and interquartile range (IQR), with whiskers extending to 1.5×IQR; jittered points represent individual sequences. The global differences were assessed using a Kruskal–Wallis test (H=159.81, P=1.09×10⁻³²), followed by Dunn’s post-hoc test with Holm–Bonferroni correction implemented using the scikit-posthocs library. Lowercase letters indicate significant differences (α=0.05): The groups sharing letters are not significantly different. The y-axis is shown on a log₁₀ scale. The number of proteins used for evaluation along with the letter of significance: Protease inhibitors from PeptipediaDB (n=25, a), fungal endopeptidases (n=75, b), Chaperons (n=255, c), plant kinases (n=44, c), fungal kinases (n=30, d), and plant endopeptidase (n=56, d)

**Figure S4.**
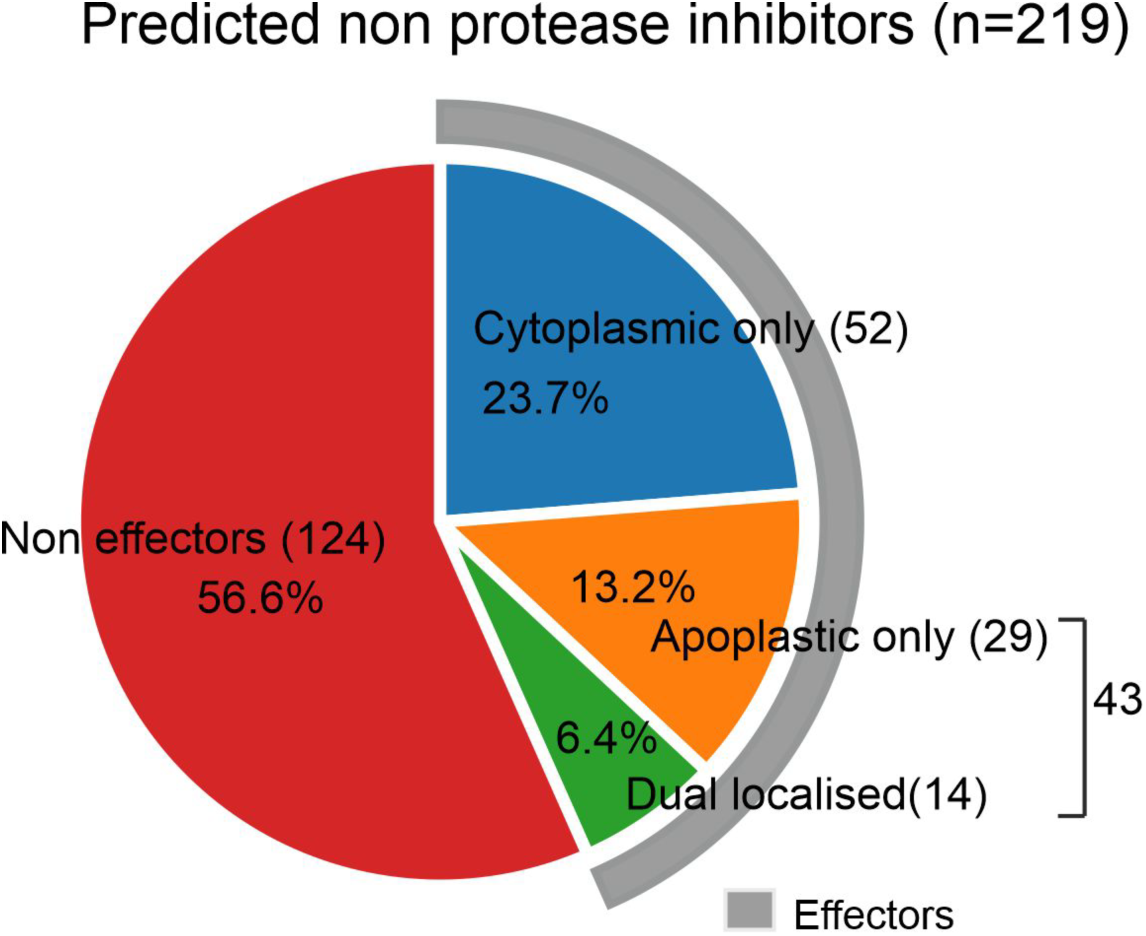
EffectorP predictions for *M. phaseolina* SSPs classified as non-protease inhibitors by the PIPES-M and PIP-BERT models at Level 1. The pie chart shows the number and percentage of proteins predicted as non-effectors, cytoplasmic effectors, apoplastic effectors, and dual-localised effectors. The outer grey highlighted ring indicates the total set of proteins predicted as effectors by EffectorP.

**Figure S5.**
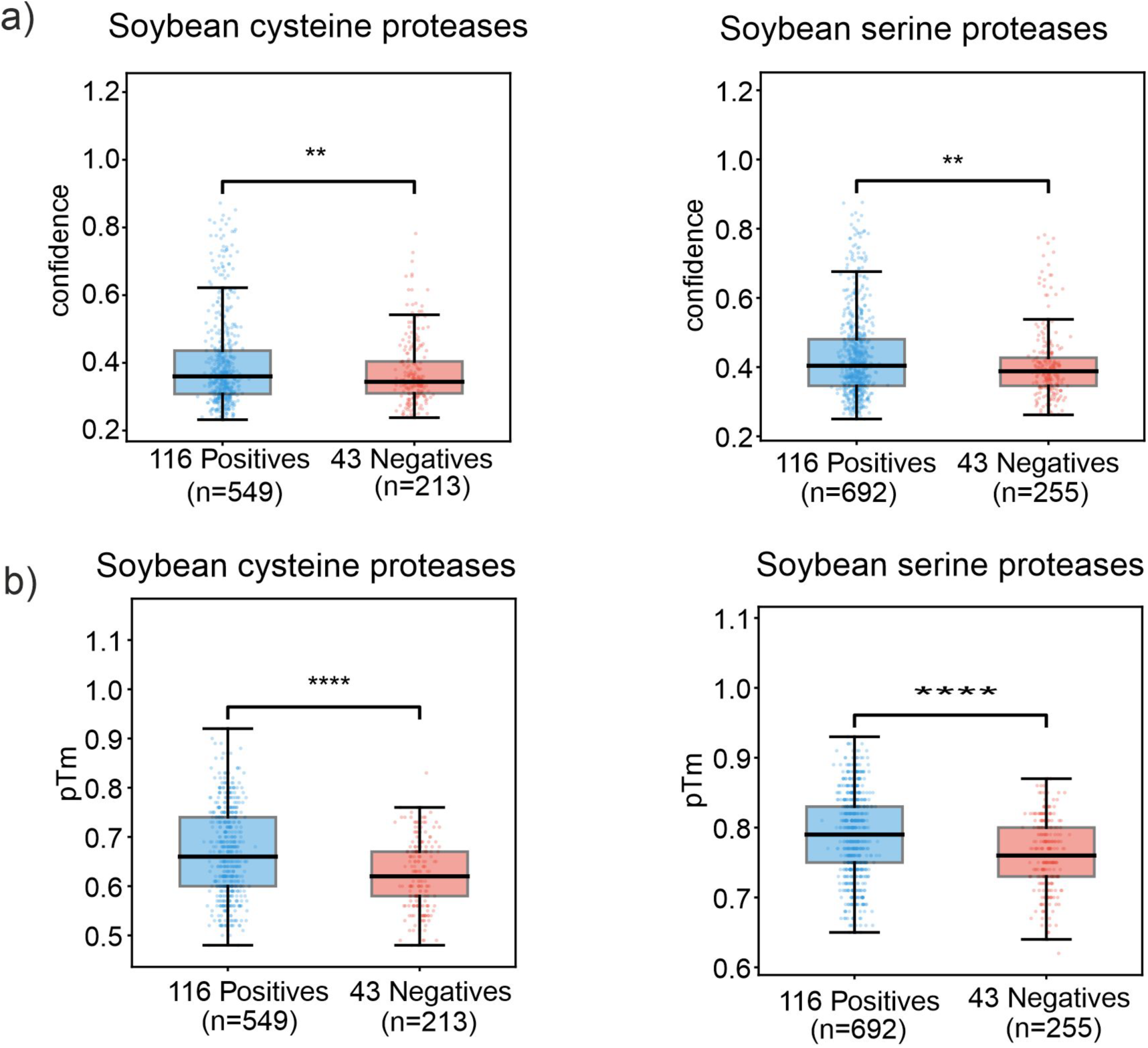
Confidence metrics for interactions between *M. phaseolina* SSPs and cysteine or serine proteases predicted by AFM. (a) Distribution of AFM confidence scores for interactions involving PLM-predicted protease inhibitors (protease inhibitors; blue) and non-protease inhibitors (non-protease inhibitors; red) across cysteine and serine proteases. (b) Distribution of pTM scores for interactions involving PLM-predicted protease inhibitors (protease inhibitors; blue) and non-protease inhibitors (non-protease inhibitors; red) across cysteine and serine proteases. The predicted protease inhibitors show significantly higher AFM confidence scores and pTM scores than non-protease inhibitors, supporting the enrichment of true inhibitor–protease interactions among the predicted protease inhibitors. Statistical significance was assessed using Welch’s t-test. Significance levels are denoted as ns (p ≥ 0.05), * (p < 0.05), ** (p < 0.01), *** (p < 0.001), and **** (p < 0.0001)

**Figure S6.**
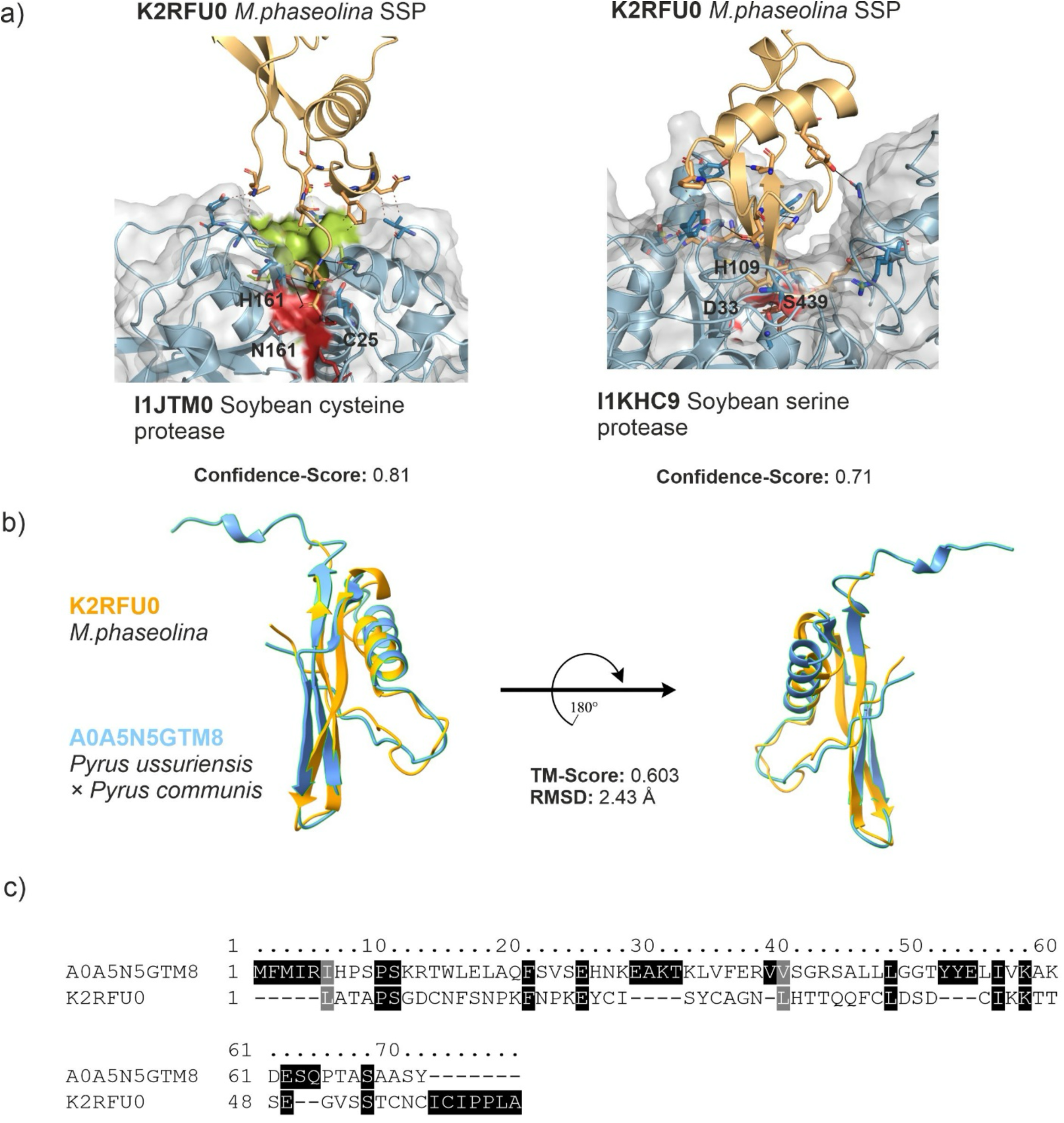
AFM predicts K2RFU0 interacts with the active site of both soybean cysteine and serine proteases. (a) AFM models of K2RFU0 (*M. phaseolina* effector) interaction with a soybean cysteine protease (UniProt Id: I1JTM0) and soybean serine protease (UniProt Id: I1KHC9). K2RFU0 is shown as a wheat coloured ribbon, and the soybean proteases are shown as grey surfaces. Interface residues are displayed as sticks and labelled. The active sites of the proteases are coloured green, and the catalytic residues are represented as red sticks. (b) Structural superposition of K2RFU0 onto experimentally resolved 3D tertiary structures in the PDB100 database was performed using Foldseek search. TM-score and RMSD values are indicated. The K2RFU0 is represented in blue, whereas crystal structures of known effectors are represented in yellow. (c) Multiple sequence alignment of K2RFU0 and A0A5N5GTM8 was generated using Clustal Omega. Conserved residues are highlighted. AFM, AlphaFold Multimer; PI, protease inhibitor; SSP, small secreted protein; PLM, protein language model; *M. phaseolina, Macrophomina phaseolina*; PyMOL, Python molecular visualisation software; PDB, Protein data bank; RMSD, root mean square deviation; TM-score, template modelling score.

